# METTL3 shapes m6A epitranscriptomic landscape for successful human placentation

**DOI:** 10.1101/2024.07.12.603294

**Authors:** Ram Parikshan Kumar, Rajnish Kumar, Avishek Ganguly, Ananya Ghosh, Soma Ray, Md. Rashedul Islam, Abhik Saha, Namrata Roy, Purbasa Dasgupta, Taylor Knowles, Asef Jawad Niloy, Courtney Marsh, Soumen Paul

## Abstract

Methyltransferase-like 3 (METTL3), the catalytic enzyme of methyltransferase complex for m6A methylation of RNA, is essential for mammalian development. However, the importance of METTL3 in human placentation remains largely unexplored. Here, we show that a fine balance of METTL3 function in trophoblast cells is essential for successful human placentation. Both loss-of and gain-in METTL3 functions are associated with adverse human pregnancies. A subset of recurrent pregnancy losses and preterm pregnancies are often associated with loss of METTL3 expression in trophoblast progenitors. In contrast, METTL3 is induced in pregnancies associated with fetal growth restriction (FGR). Our loss of function analyses showed that METTL3 is essential for the maintenance of human TSC self-renewal and their differentiation to extravillous trophoblast cells (EVTs). In contrast, loss of METTL3 in human TSCs promotes syncytiotrophoblast (STB) development. Global analyses of RNA m6A modification and METTL3-RNA interaction in human TSCs showed that METTL3 regulates m6A modifications on the mRNA molecules of critical trophoblast regulators, including *GATA2, GATA3, TEAD1, TEAD4, WWTR1, YAP1, TFAP2C* and *ASCL2*, and loss of METTL3 leads to depletion of mRNA molecules of these critical regulators. Importantly, conditional deletion of *Mettl3* in trophoblast progenitors of an early post-implantation mouse embryo also leads to arrested self-renewal. Hence, our findings indicate that METLL3 is a conserved epitranscriptomic governor in trophoblast progenitors and ensures successful placentation by regulating their self-renewal and dictating their differentiation fate.

## Introduction

In mammals, placentation is a remarkable adaptive response for successful reproduction. During the development of the placenta, trophoblast/stem progenitor cells (TSPC) orchestrate the initiation and differentiation of extra-embryonic trophoblast lineages ^1^. These differentiated cells have evolved distinct temporal and spatial characteristics to fulfill specific functions at the site of implantation, exchange of nutrients, oxygen, metabolites, and the establishment of maternal-fetal interface ^2–4^. Improper development of the trophoblast cell lineages or aberrations in trophoblast cell function have been associated with adverse pregnancy outcomes leading to failure of embryo implantation, the Great Obstetric Syndrome (e.g., intrauterine fetal growth restriction (IUGR), preeclampsia (PE), preterm birth or extreme preterm birth) and intrauterine lethality leading to recurrent pregnancy loss (RPL) ^5–8^ or increasing the risk for the development of severe disorders in later life such cardiovascular, metabolic/obesity, neuropsychiatric disease and type 2 diabetes ^9,10^. Each of these conditions poses risks to both maternal and fetal well-being, imposing considerable health and socioeconomic burdens. Therefore, unraveling the molecular mechanisms behind these pregnancy complications is crucial from both clinical and economic standpoints.

A developing first-trimester human placenta contains two types of villi; (i) Floating villi, which float into intervillous space and (ii) anchoring villi, which attach to the maternal endometrium ^11^. A floating villous contains two different layers of trophoblast cells; (i) the cytotrophoblast (CTB) progenitors, close to the stroma and (ii) the post-mitotic STB layer overlaying the CTBs. The STBs establish the main uterine-placental interface for nutrient and gas exchange between the mother and the developing fetus. In the anchoring villi, CTBs establish a column of proliferating CTB progenitors, known as column CTBs (CCTs). CCTs differentiate into migratory invasive EVT cells, which invade into the maternal uterine compartment. A subset of EVTs, which invade the uterine compartment, remodel the uterine artery for increased blood flow at the uterine-placental interface for adequate nutrient supply to the developing fetus. These EVTs are known as endovascular EVTs ^12–14^. The remaining invasive EVTs within the uterine interstitium comprise the interstitial EVTs ^14^, which interact with uterine cells for adaptation of the maternal immune system and physiology to the developing placenta. CTB progenitors are the source of STBs and EVTs and proper maintenance of CTB self-renewal and their coordinated differentiation to STBs and EVTs are essential for the initiation and maintenance of placental structure and functions throughout gestation ^3,4,15–19^. The maintenance of self-renewal in CTBs and establishment of STB *vs*. EVT differentiation potential is a highly dynamic process and relies on the molecular mechanisms that fine-tune the gene expression programs in different CTB progenitor subpopulations ^20–31^. However, the importance of RNA epigenetic (epitranscriptomic) regulations for the maintenance of CTB self-renewing state and induction of STB and EVT differentiation fates remains poorly understood.

The epitranscriptomic regulation *via* N^6^-methyladenosine (m^6^A) in eukaryotes plays a vital role in diverse physiological and pathological conditions ^32–34^. The m6A modification is achieved by the core catalytic subunit METTL3 of the methyltransferase complexes (MTC complexes) in the nucleus ^35–37^ and can be specifically blocked using pharmacological catalytic inhibitor of METTL3, such as STM2457 ^38^. METTL3 and its heterodimeric partner METTL14 are evolutionarily conserved catalytic subunits of MTC complexes and are essential for early mammalian development as either *Mettl3 or Mettl14* global knockout mice die during early post implantation stages ^35,39^. It has further been demonstrated that METTL3 is critical for embryo implantation and decidualization ^40^. Recently, it has also been reported that in preeclampsia, a human pregnancy associated pathological condition, METTL3 is upregulated which may contribute to trophoblast dysfunction in preeclampsia *via* aberrant m6A modification ^41–43^.

Here, we tested the importance of METTL3 in trophoblast development. We studied human TSCs established earlier ^44^ as well as human TSCs, which we established from pregnancies associated with idiopathic RPLs. In addition, we studied placental samples from human pregnancies associated with FGR and preterm birth. Our findings reveal that improper expression levels of METTL3 have detrimental effects on both the self-renewal and the differentiation potential of human trophoblast progenitor cells. METTL3-deficient human TSCs spontaneously differentiate into STBs and fail to differentiate into EVTs. Using RNA CUT& RUN-sequencing to identify global m6A modification and METTL3-fRIP-seq we demonstrate that transcripts of several crucial genes necessary for the self-renewal and EVT differentiation of human TSCs undergo METTL3-dependent m6A modification. Using conditional *Mettl3-* KO mouse model we show that METTL3 is essential for the self-renewal of trophoblast progenitors of the developing mouse placenta. Our findings establish the METTL3-mediated m6A modification underlies proper trophoblast development during human placentation.

## Results

### METTL3 expression is conserved in the placental trophoblast progenitors of developing mouse, rat, and human placenta

Development of the trophoblast cell lineage begins with the specification of the trophectoderm (TE) during morula to blastocyst transition. TE cells are specialized for implantation and interaction with the maternal environment. In rodents, after embryo implantation, multipotent trophoblast stem and progenitor cells (TSPCs) arise from the TE. TSPCs undergo extensive proliferation and differentiation to develop the placental primordium, consisting of the extraembryonic ectoderm (ExE)/ectoplacental cone (EPC) and the chorionic ectoderm ^1^. To understand the importance of METTL3 and m6A in trophoblast development during placentation, first we tested the expression of METLL3 in the mouse trophectoderm (TE) and in TSPCs of early post implantation mouse embryo. We found that METTL3 protein is robustly expressed in the nuclei of TE cells (**Fig. 1A**) and in TSPCs within the ExE/EPC region of an embryonic day 7.0 (E7.0) mouse embryo (**Fig. 1B**).

**Fig. 1.**
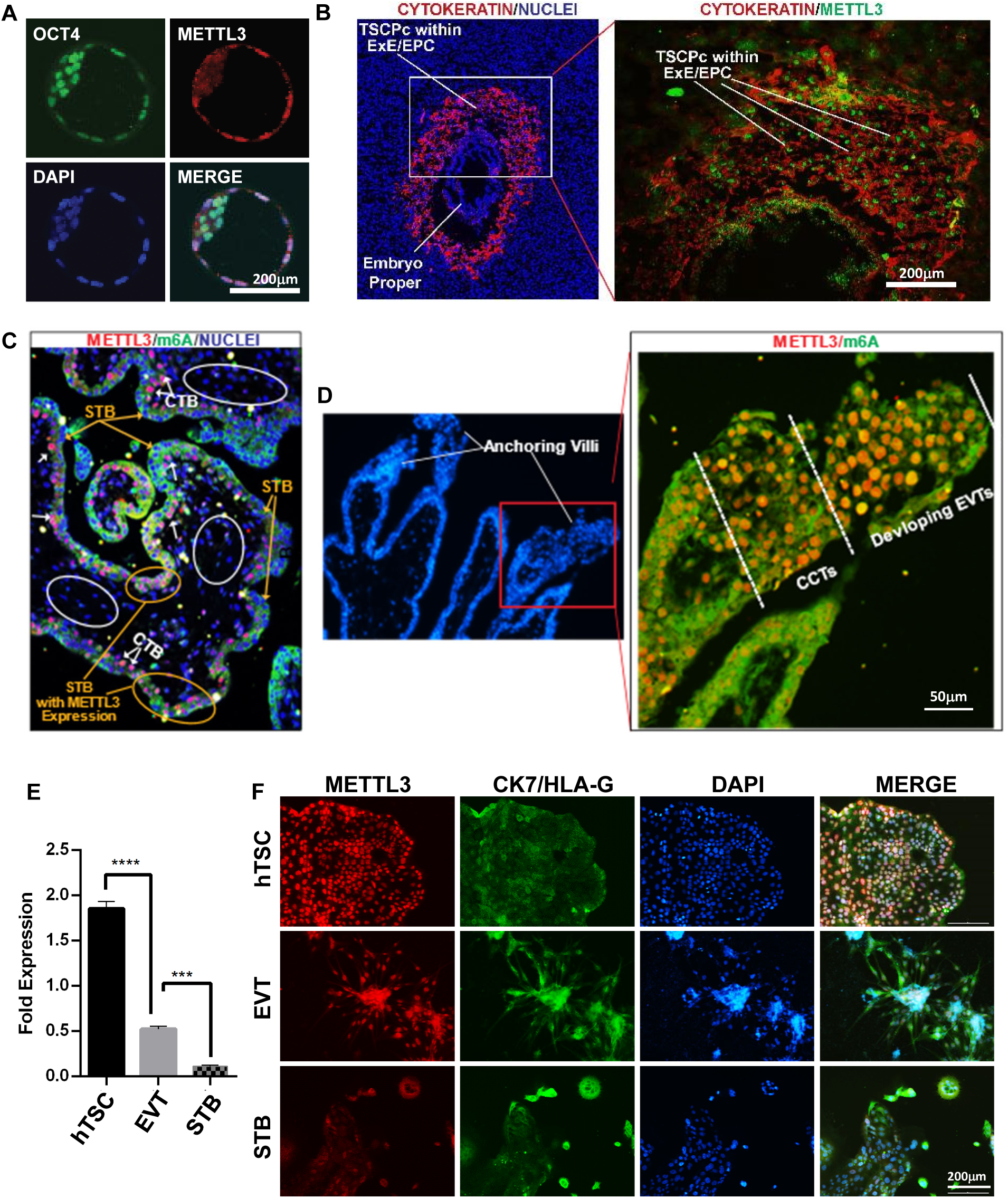
METTL3 expression is conserved in rodent and human trophoblast progenitors. (A) Immunohistochemistry showing expression of METTL3 (red) and OCT4 (green) in the mouse blastocyst. (B) An embryonic day 7.0 mouse extraembryonic ectoderm (ExE)/Ectoplacental cone (EPC) region showing METTL3 (green) and Cytokeratin (PanCK, red) expressions in TSPCs. (C) Histological section of a human first-trimester (week 6) placenta showing METTL3 expression and m6A modification in the floating villi. METTL3 expression is mostly confined to CTBs (white arrows), whereas STB nuclei show a mosaic expression pattern. Some STB nuclei express METTL3 (yellow bars), whereas expression is repressed in other STB nuclei (yellow arrows) and most of the stromal cells (white ellipses). All the cells expressing METTL3 are enriched for m6A modification. (D) Immunofluorescence showing METTL3 expression in an anchoring villi of a first-trimester human placenta (6 week). METTL3 is highly expressed in both CCTs and developing EVTs. All the cells expressing METTL3 are also enriched for m6A modification. (E) RT-qPCR data showing expression of *METTL3* in hTSCs, differentiated EVTs and STBs. (n=3 independent experiments, p≤0.0001). (F) Immunohistochemistry showing expression of METTL3 protein levels in the hTSC, EVTs and STBs. It is important to note that METTL3 is only detected in some of the nuclei within the STB and lower than the hTSC or EVT. Scale bar 200μm.

To begin to understand the functional importance of METTL3 in human trophoblast development, we tested METTL3 expression and m6A modification in trophoblast cells of a first-trimester human placenta. As mentioned earlier, in a first-trimester human placenta floating villi contain CTB progenitors and the post-mitotic STB layer overlaying the CTBs, whereas the anchoring villi contain CCTs and emerging EVTs that differentiate from CCTs. Immunofluorescence analyses revealed that in the floating villi the m6A modification is enriched in both CTBs and STBs (**Fig. 1C**, green). We noticed that METTL3 expression is mostly confined to the CTB progenitors (**Fig. 1C**, white arrows). In contrast to CTBs, METTL3 expression is repressed in the majority of STB nuclei (**Fig. 1C**, yellow arrows). However, we noticed METTL3 protein expression in some patches of STB nuclei (**Fig. 1C**, yellow ellipses). We also noticed that most of the stromal cells lack METTL3 expression as well as m6A modification (**Fig. 1C**, white ellipses). In the anchoring villi, both CCTs and developing EVTs abundantly express METTL3 and are enriched with m6A modification (**Fig. 1D**.). Taken together, we concluded that in a developing first-trimester human placenta METTL3 expression is confined within the CTB progenitors and in developing EVT cells. In contrast, the STB differentiation in floating villi is associated with suppression of METTL3 expression. We also concluded that patches of METTL3 expressing STBs represent areas where CTB nuclei may be freshly fused to form nascent STB layers.

Next, we tested expression of METTL3 in CTB-derived hTSCs ^20,44^. Akin to the expression profile in primary trophoblast cells of human first-trimester placentae, undifferentiated hTSCs and hTSC-derived EVT cells exhibited elevated levels of METTL3 RNA and protein expressions compared to STBs (**Fig. 1E-F**). We also confirmed the expression patterns of *METTL3* in primary trophoblast cells at a single-cell resolution by reanalyzing recently published single cell RNAseq (scRNAseq) data from first-trimester placentae ^20,21^. We noticed that *METTL3* and the member of MTC complex *METTL14* are expressed in all cell populations of first-trimester placentae except within the mature STB cell cluster (Supp Fig. S1A).

Similarly, re-analyses of recently published scRNAseq data in hTSCs ^45^ also confirmed *METTL3* expression pattern in undifferentiated hTSCs and upon their differentiation in EVTs and STBs (Supp Fig. S1B). As METTL3 is highly expressed in invasive EVT population, we tested whether METTL3 expression is conserved in invasive trophoblast cells in other species. We used rat as an experimental model as it shows deep trophoblast invasion at the uterine-placental interface ^46^. We confirmed that like human EVTs, rat invasive trophoblast cells highly express METLL3 (Supp Fig. S1C) The conserved expression pattern of METTL3 prompted us to hypothesize that METLL3 is important to orchestrate gene expression in human trophoblast progenitors to dictate their self-renewal and differentiation fate and impaired METTL3 function could be associated with adverse pregnancy outcome. Therefore, we tested whether human pathological pregnancies are associated with impairment of METTL3 expression in trophoblast cells.

### Imbalanced METTL3 expression levels in human trophoblast progenitors are associated with pregnancy-related complications

In human, developmental abnormalities during placentation, including defect in trophoblast development, are associated with recurrent pregnancy loss (RPL) or pregnancy associated complications such as preterm birth, intrauterine growth restriction (IUGR) and preeclampsia. These abnormalities disrupt normal placental development and function, impacting the health of both the mother and the fetus. To investigate the biological significance of METTL3 in association with human pregnancy related complications, we checked *METTL3* mRNA expression in placentae associated with IUGR and preterm birth. We observed a significant downregulation of *METTL3* transcript levels in preterm placentae compared to term placentae (**Fig. 2A**). Intriguingly, in contrast to the preterm pregnancies, *METTL3* transcript levels in IUGR placentae exhibited a substantial upregulation (**Fig. 2A**). Analyses of METTL3 protein levels also confirmed downregulation of METTL3 in preterm placentae compared to term placentae (**Fig. 2B**) and significant upregulation of METTL3 in IUGR placentae (**Fig 2B**).

**Fig. 2.**
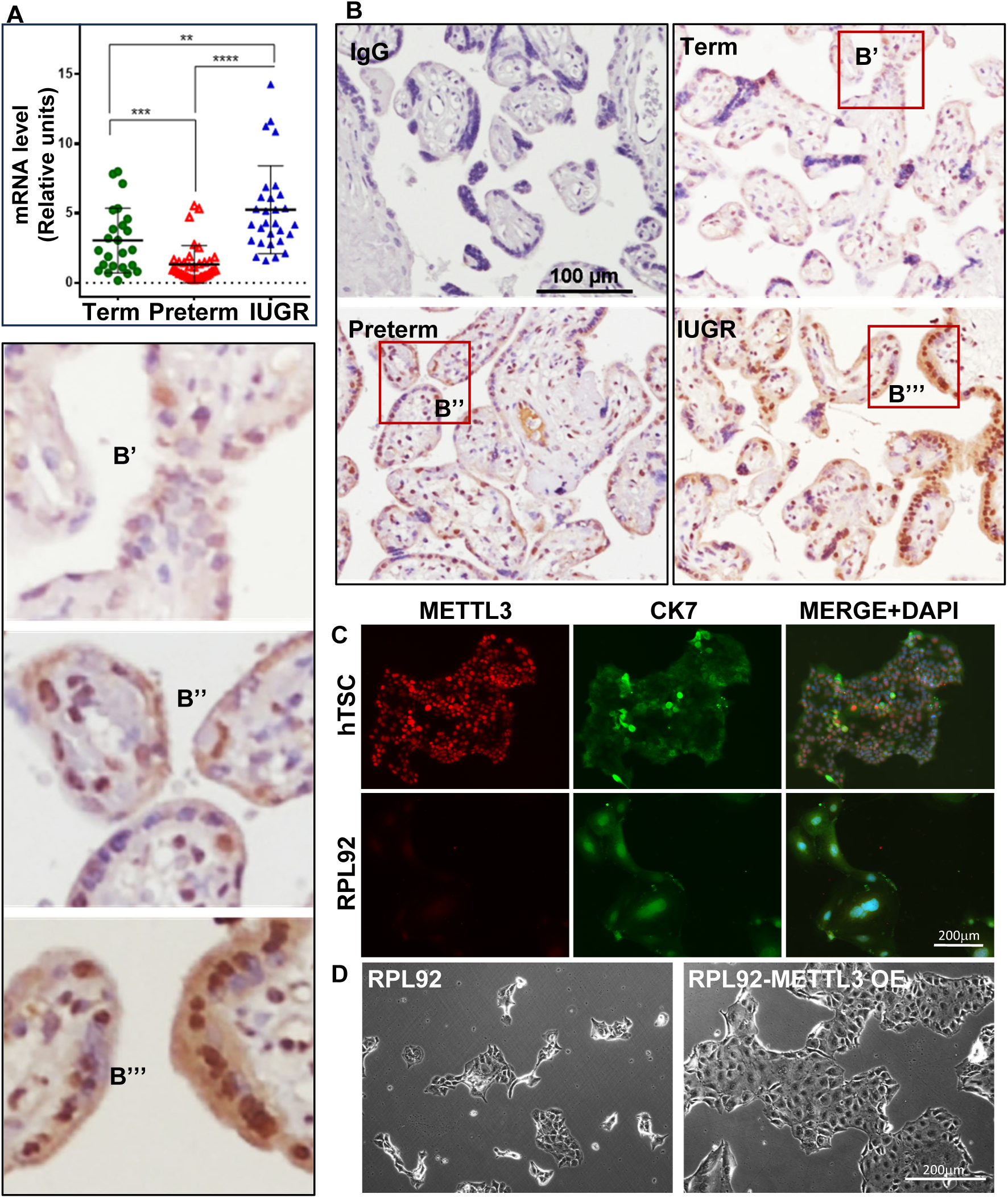
Imbalanced METTL3 expression in human placenta is associated with pregnancy related complications. (A) RT-qPCR analyses of *METTL3* mRNA expression in placental tissues from pregnancies with term control (gestational age ≥38 weeks), preterm-birth (≤36 weeks) and IUGR. [Error bars, mean ± SEM, * Indicates significant change (*\*\** < 0.01, \**\*\** < 0.001, \*\**\*\** < 0.0001)]. (B) Representative immunostained images depicting METTL3 expression in placentae from pregnancies associated with term, preterm, and IUGR births reveal noteworthy differences. METTL3 expression appears lower compared to term controls in preterm cases, while it is upregulated in IUGR [(B’-B’’’ inset enlarged (scale 100μm)]. (C) Representative immunostained images of control CT27 hTSC (upper panel) and RPL92 hTSC (lower panel) showing METTL3 (red) and Cytokeratin (CK7, green) expressions (scale 200μm). Note severe loss of METTL3 expression in RPL92 hTSCs. (D) Rescue of cell proliferation in RPL92 hTSC lines upon ectopic rescue of METTL3 expression. Representative micrograph shows cell colonies upon culturing for 72h. (scale 200μm).

To further understand the correlation of METTL3 expression in trophoblast progenitors with adverse human pregnancies, we tested METTL3 expression in the context of idiopathic RPL. Earlier, we reported that a subset of idiopathic RPLs is associated with major defects in placental villi formation, characterized by defective formation of the CTB/STB bilayer ^21^. We also isolated CTBs and established idiopathic RPL patient-specific hTSC lines (RPL-TSCs) to understand trophoblast intrinsic causes that might lead to idiopathic RPL. Upon careful analysis of 57 RPL-TSC lines, we identified 8 RPL-TSC lines, showed major defects in proliferation in which METTL3 protein expression was extremely low or was undetectable (**Fig. 2C**, **Supp Fig. S3**). To further assess METTL3 expression level for sustaining the proliferation potential of hTSCs, we selected an RPL-TSC line, RPL92, which exhibited undetectable levels of METTL3, and ectopically expressed *METTL3*. Remarkably, the ectopic expression of METTL3 effectively rescued the proliferation potential of RPL-TSC line 92 (**Fig. 2D**). Collectively, our studies on pathological pregnancies strongly indicated that a fine balance in METTL3 function is required for human trophoblast development and defect in METTL3 function might lead to defective placentation due to impaired trophoblast progenitor maintenance.

### METTL3 regulates the self-renewal potential of hTSCs and dictates their differentiation fate

To further investigate the functional importance of METTL3 in human trophoblast development and function, we depleted *METTL3* in hTSCs by RNAi using lentiviral-mediated transduction. Initially, we used constitutively expressing shRNAs against *METTL3*, which resulted in a >80% reduction in *METTL3* transcript levels, leading to the complete loss of TSC stem-state colony morphology. Consequently, we opted to employ doxycycline-inducible shRNA (*tetOshMETTL3)* with the same sequence to conditionally deplete *METTL3* in hTSCs. hTSCs expressing *tetOshMETTL3* were continuously treated with doxycycline to assess *METTL3* knockdown. We noticed that doxycycline treatment for a duration of 4 days resulted in ∼80% reduction in *METTL3* transcript levels and undetectable levels of METTL3 protein expressions in hTSCs (**Fig. 3A, B C**). Under this experimental condition, we observed a robust reduction in m6A-modified RNA (**Fig. 3D**) and hTSC proliferation, assessed *via* BrdU incorporation (**Fig. 3E-F**) and the number of mitotic nuclei (**Fig. 3G, I**). Interestingly, upon METTL3 depletion, another notable observation emerged: the surviving METTL3-depleted hTSCs frequently adopted a STB-like morphology, characterized by multinucleation (**Fig. 3G, J**) and significantly elevated expression of the STB marker hCGβ (**Fig. 3K** and **Supp Fig. S4A**) while maintained in stem state culture condition. In contrast, the *METTL3*-depleted cells did not exhibit induction of HLA-G, a marker for EVT differentiation (**Supp Fig. S4B**).

**Fig. 3.**
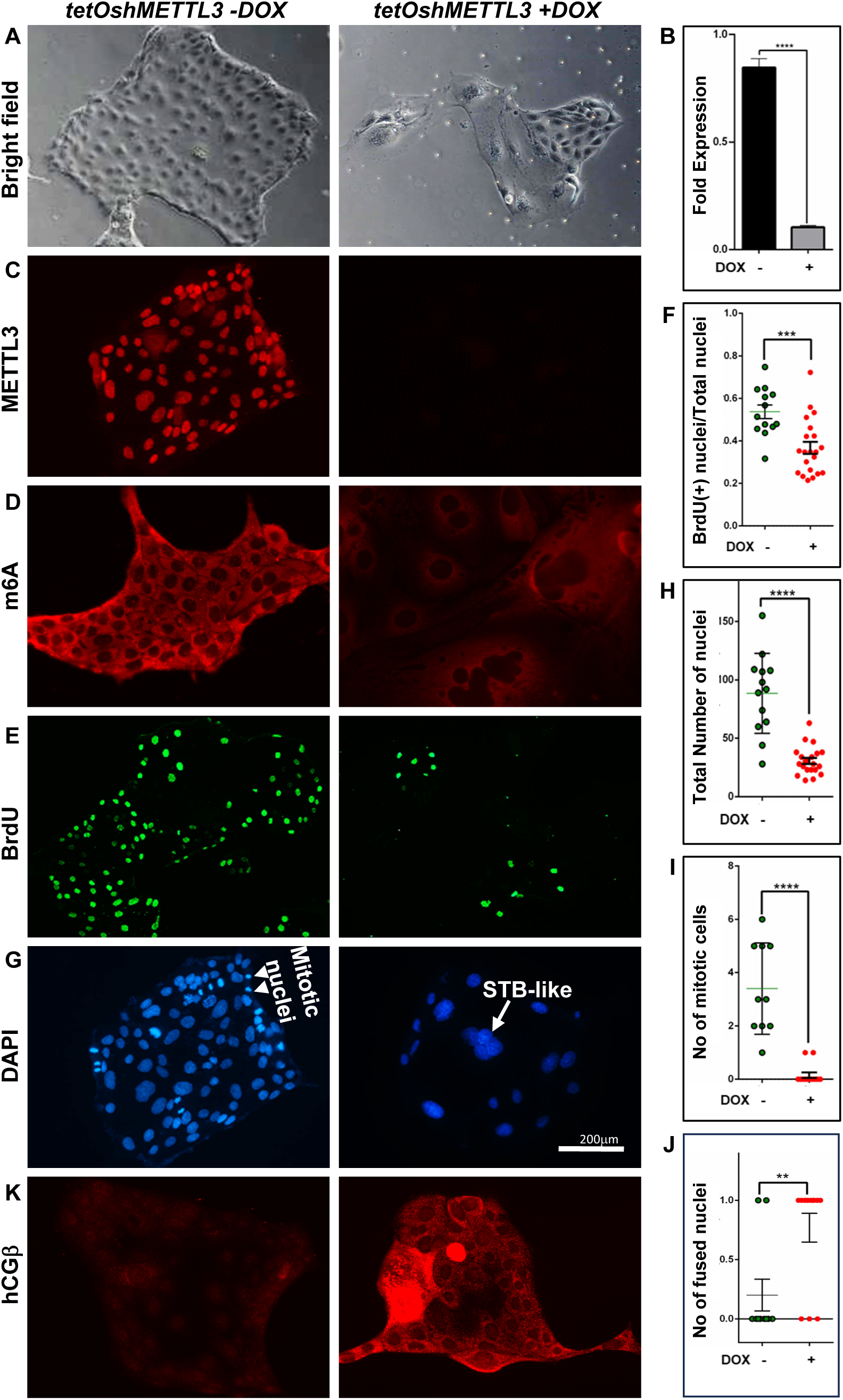
hTSC lacking METTL3 results in impaired self-renewal potential and concomitantly differentiate into STBs. (A) Representative micrograph showing shRNA mediated knockdown of *METTL3* in CT27 hTSCs reduced cell proliferation, (B) RT-qPCR showing *METTL3* mRNA knockdown efficiency (n=3 independent experiments, p<0.0001). (C) Representative immunostained colonies of hTSC showing METTL3 knockdown efficiency at protein level. (D) Representative immunostained colonies of CT27 hTSC showing loss of METTL3 reduces global m6A modification. (E) Representative immunostained colonies of CT27 hTSCs show BrdU incorporation in control and *METTL3* depleted hTSC when cultured in stem-state culture condition at 72 hours. (F) BrdU positive nuclei quantitation showing loss of *METTL3* abrogate rate of hTSC proliferation (n=3 independent experiments, p<0.001). (G) Representative DAPI images depicting hTSC nuclei used to quantitate for data presented in panels (H-J). (H) Plot shows total number of nuclei/10x microscopic fields (representative of intact cells) is severely reduced upon *METTL3* depletion (n=3 independent experiments, p<0.0001). (I) Quantitation of mitotic nuclei/20x microscopic field in control and *METTL3*-depleted CT27 hTSCs upon culturing for 72h in stem state culture condition (n=3 independent experiments, p<0.001). (J) Quantitation of fused nuclei//20x microscopic field (as a measure of STB differentiation) in control and METTL3-depleted CT27 hTSCs upon culturing for 72h in stem state culture condition (n=3 independent experiments, p<0.01). (K) Representative CT27 hTSC colonies, maintained at stem state culture condition, showing hCGβ induction upon loss of METTL3. Scale 200μm.

The self-renewing ability of hTSCs and CTB progenitors can also be assessed by their ability to generate 3D-trophoblast organoids. The trophoblast organoids grow in an inside-out pattern, in which the self-renewing hTSCs/CTBs grow as an outer layer, whereas the cells inside the 3D-organid undergo STB differentiation ^20,21,47–49^. We tested the self-renewal efficiency of *METTL3*KD hTSC by assessing their ability to form three-dimensional trophoblast organoids (TSC 3D organoids) with prolonged culture (8-10 days). Unlike the control hTSCs, *METTL3*KD human TSCs showed severe impairment in organoid formation (**Fig. 4A**). To further assess the self-renewing ability primary 3D hTSC organoids were dissociated and replated to form the secondary organoids. In contrast to the control hTSC organoids, *METTL3*KD human TSCs failed to develop secondary organoids.

**Fig. 4.**
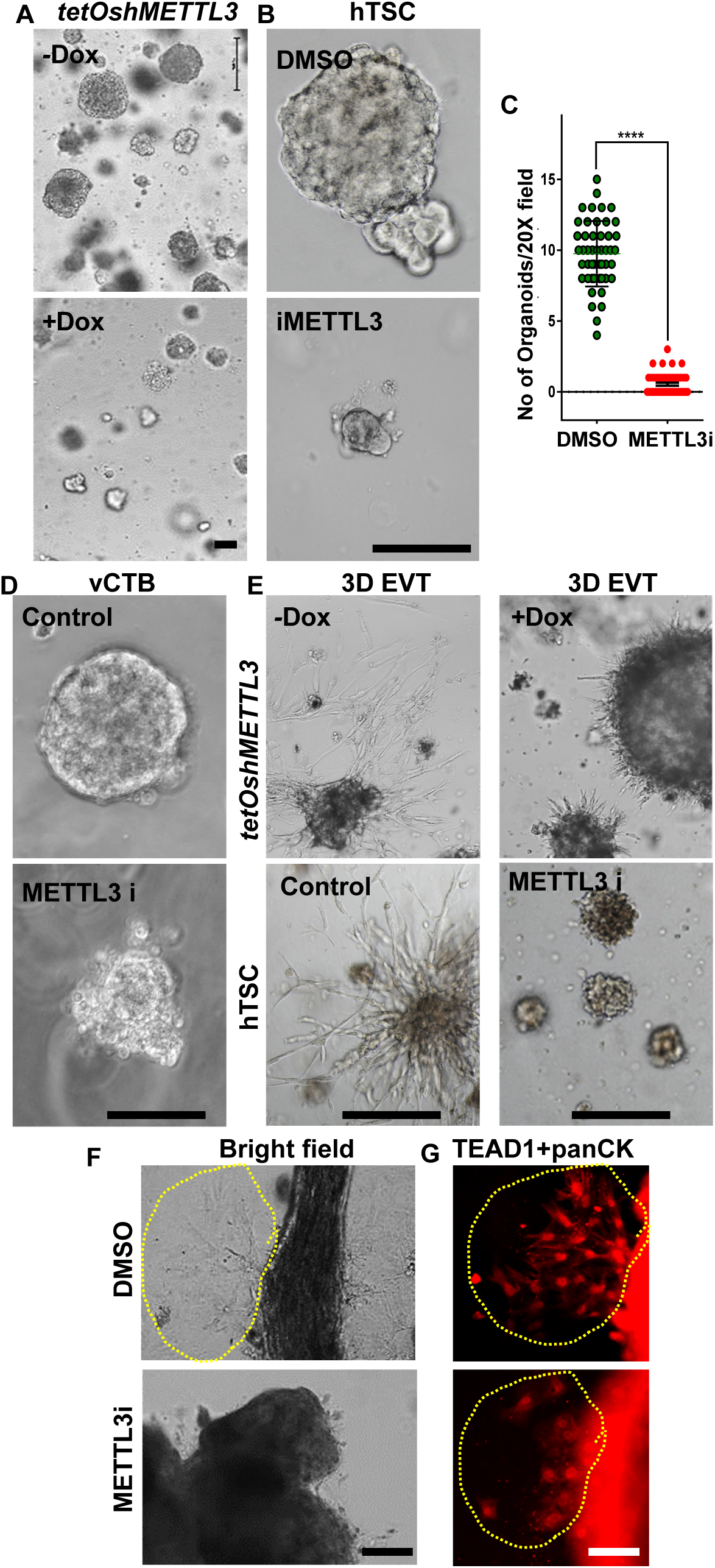
Loss of METTL3 in hTSCs impairs both self-renewing 3D organoid formation and EVT differentiation. (A) Representative micrograph showing depletion of METTL3 in hTSCs inhibit 3D hTSC organoid formation. (B) Representative micrographs showing that inhibition of METTL3 with METTL3 inhibitor STM2457 (METTL3i) in hTSCs inhibits 3D organoid formation. (C) Quantitation of total number of 3D organoids for the panel B (n= 3 independent experiments, p<0.0001). (D) Representative micrograph showing that inhibition of METTL3 in primary villus cytotrophoblasts (vCTBs) inhibit organoid formation. It is important to note that METTL3i-treated primary vCTB organoids completely fail to form secondary organoids upon passaging. (E) Representative phase contrast images showing 3D hTSC organoids in a culture condition that promotes EVT differentiation. Control TSCs organoids readily developed EVTs migrating out from the center of the organoid (Left panel). The EVT development was strongly impaired from hTSC organoids with METTL3-depletion (upper panel) or treated with METTL3i (bottom panel). (F) Representative phase contrast images showing that EVT development is impaired from METTL3i-treated first-trimester human placental explants when cultured on Matrigel in a culture condition that promotes EVT differentiation. (G) Representative immunostained images of TEAD1 (EVT marker) showing emergence of EVT from the control placental explants and inhibition of EVT development upon treatment with METTL3i (scale 200μm).

We also tested the importance of METTL3 for self-renewal of primary villous CTBs, isolated from first-trimester human placentae. To that end, we leveraged a highly selective catalytic METTL3 inhibitor, STM2457 (henceforth mentioned as METTL3i), which selectively inhibits METTL3-mediated m6A modification ^38^. Interestingly, hTSCs and primary CTBs, treated with METTL3i (10μM, treated from day 2-6), exhibited significant impairment in the trophoblast organoid formation. In contrast to the control hTSCs and CTBs (treated with DMSO), the METTL3i treated hTSCs and CTBs formed significantly smaller organoids (**Fig. 4B-D**).

As METTL3 is highly expressed in CCTs and developing EVTs within and anchoring villi, we tested the importance of METTL3 in EVT development using three different experimental approaches. First, we tested EVT differentiation efficiency of *METTL3*-KD hTSCs in 3D-organoid culture conditions. We found that loss of METTL3 in hTSCs strongly impaired EVT differentiation potential in 3D-hTSC-derived organoid model. EVT emergence was readily observed from control hTSC-organoids. However, loss of METTL3 through doxycycline-inducible RNAi as well as inhibition of METTL3 with METTL3i abolished EVT development from 3D-hTSC organoids (**Fig. 4E**). Finally, we assessed EVT emergence from human first-trimester placental explants in the presence and absence of METTL3i. Consistent with our observations in hTSC 3D organoids, the inhibition of METTL3 with METTL3i impeded EVT emergence from first-trimester placental explants (**Fig. 4F-G**).

Collectively, from our studies in human TSCs, primary CTBs and placental explants we posit that METTL3-mediated m6A modification is essential to maintain self-renewal ability in CTB progenitors. Our findings also indicate that, during human placentation, in the floating villi, METTL3 functions as a gatekeeper in CTBs to prevent premature adaptation of STB fate. In contrast, within anchoring villi METTL3 function is essential to adapt EVT differentiation fate of CCTs.

### METTL3 mediated m6A modification is essential for stochiometric balance of key transcripts, essential for human trophoblast development

To understand how METTL3 governs the gene expression program to sustain stemness in hTSCs, we performed unbiased gene expression analysis through RNA sequencing (RNA-seq). Comparison of RNA-seq data win control hTSCs *vs*. *METTL3KD* hTSC identified significantly altered expression of 7453 genes (3661 upregulated and 3792 downregulated (foldchange >1.5) (**Fig. 5A-B**, **Dataset S1, sheet1**). We used EnrichR ^50^ to assess association of METTL3-regulated differentially expressed genes (DEGs) in hTSCs. METTL3-regulated DEGs show strong association with embryonic development, perinatal lethality, and postnatal growth retardation (**Supp Fig. S5A).** Interestingly, loss of METTL3 also strongly altered genes that are associated with WNT and TGFβ signaling pathways, which are key signaling components to regulate CTB progenitor state and EVT differentiation (**Supp Fig. S5B)** as well as cytoplasmic translation and cellular respiration **(Supp Fig. S5C)**. We also identified that METTL3 is important to maintain mRNA levels of many hTSC growth regulators, identified by Dong *et al.*,^51^. Many of these growth regulators are either downregulated or upregulated in *METTL3KD* hTSCs (**Dataset S1, sheet2**), indicating that METTL3 is important to maintain proper transcriptional stoichiometry of these important regulators in hTSCs. Analyses of our single-cell RNA-seq data ^21^ from first-trimester human placenta indicated that many of these METTL3-regulated transcripts are highly induced in either CTB progenitors at their stem state or when they are undergoing EVT differentiation (**Fig. 5D**, **Supp Fig. S6**). Analyses of gene expression data from Okae *et al.,* study ^44^ further confirmed that these genes are selectively induced in CTBs and/or EVTs but are downregulated in STBs. PlacentaCellEnrich ^52^ analyses of METTL3-regulated DEGs revealed that upregulated genes in *METTL3*KD hTSCs have very strong association with STB differentiation (**Supp Fig. S7A**), which we further confirmed by comparing RNA-seq data with Shimizu *et al.*, study ^23^, which identified genes that are strongly associated with STB differentiation (STB hub genes, **Dataset S1, sheet3**)). We confirmed that many of those STB hub genes are upregulated in *METTL3*-KD hTSCs (**Supp Fig. S7B**). In contrast, genes that were downregulated in *METTL3*KD hTSCs showed strong association with EVT development or normally expressed in non-trophoblast cells of a human placenta (**Supp Fig. S7C**). Taken together, our unbiased gene expression analyses aligned with our phenotypic observation that the loss of METTL3 in hTSCs leads to defective self-renewal and EVT differentiation and promotes STB differentiation.

**Fig. 5.**
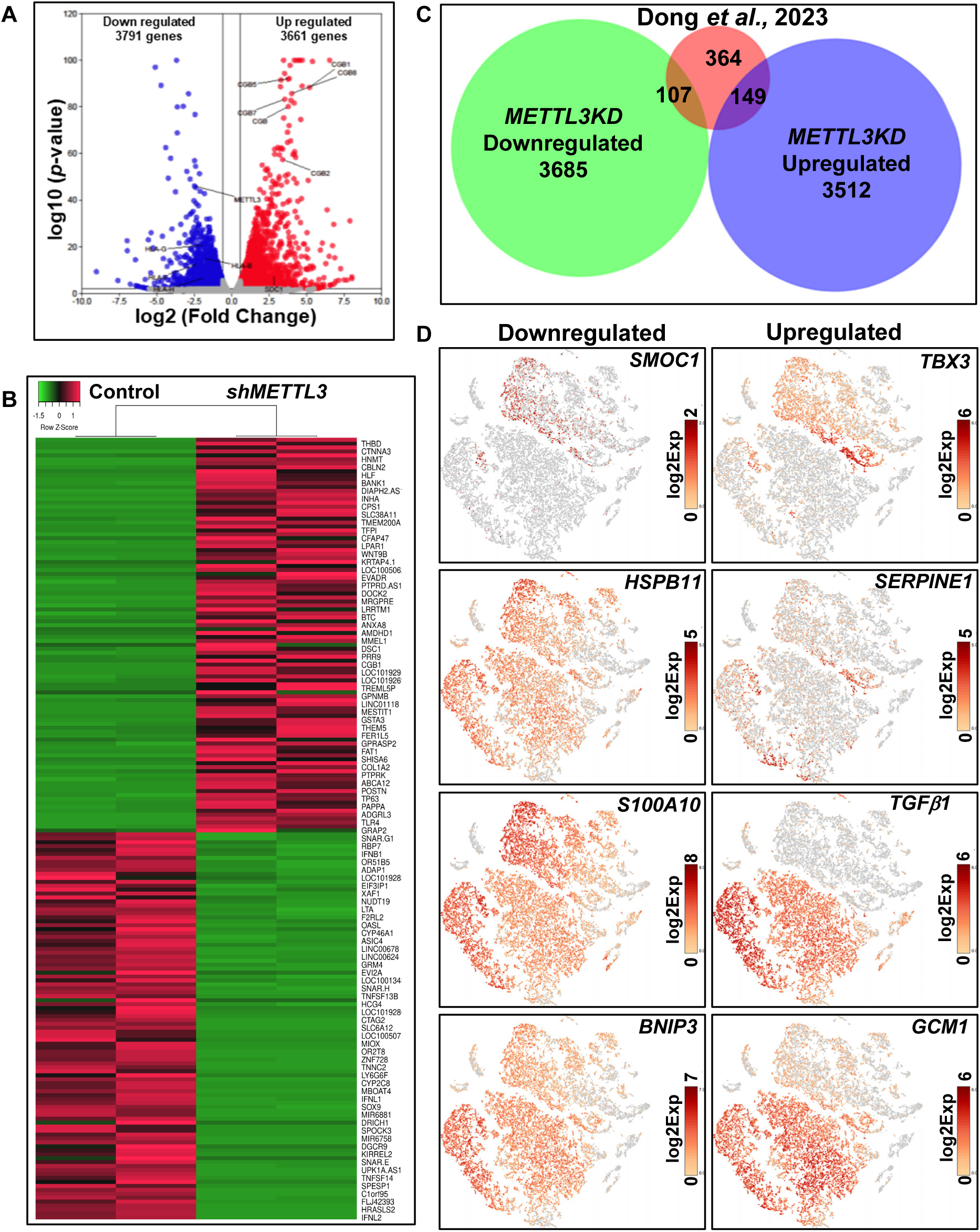
METTL3 governs trophoblast stem state gene expression dynamics. (A) Volcano plot showing DEGs (fold change >1.5 and p value < 0.05) between control and *METTL3KD* hTSC (red: up-regulated, blue: downregulated). Some of the differentially expressed genes are indicated. (B) Heat map of top 100 up-regulated and down-regulated (fold change >1.5 and *p* value < 0.05) genes between control and METTL3KD hTSC (red: up-regulated, green: downregulated genes). (C) Venn diagram showing the METTL3 regulated DEGs. The smallest circle indicates number of METTL3-regulated hTSC growth regulators, identified by Dong *et al*., 2023 study (Ref. 22). (D) t-SNE plots showing the differential mRNA expression patterns of METTL3 regulated key hTSC regulators in single-cell clusters (outlined in Supp Fig S1), obtained by scRNA-seq analyses in first-trimester human placentae (Data was rederived from Saha *et al.,* 2020 study (Ref 21)).

METTL3 complexes with METTL14 ^53,54^ and often recognizes specific RNA sequences, commonly the most preferred CUGCAG motif, for subsequent modification of adenosine to m6A ^55^. Interestingly, METTL3 also binds chromatin and regulates chromatin accessibility and transcription either *via* m6A modifications on chromosome-associated regulatory RNAs ^35^ or *via* promoting histone modifications ^56^. Alteration of m6A modification in several candidate genes has been implicated in pathological pregnancies, including IUGR and Preeclampsia ^41–43^. However, global m6A modification and interrelationship of m6A modification with gene expression program in human trophoblast progenitors are yet to be defined. Thus, to gain mechanistic insights about METTL3-mediated orchestration of hTSC transcriptome, we performed three experiments; (i) we captured global m6A modification on RNA transcripts in hTSCs *via* m6A RNA CUT&RUN **(Supp Fig. S8A)**, (ii) we performed METTL3-fRIP to capture METTL3 target RNAs in hTSCs **(Supp Fig. S8B)**, and (iii) we performed CUT&RUN to identify METTL3 occupied chromatin regions in hTSCs (**Fig. 7B**).

RNA CUT&RUN analyses identified 8008 m6A peaks in control hTSCs and GREAT analyses assigned 8008 m6A peaks in control hTSCs (**Dataset S2, sheet1**). In contrast, RNA CUT&RUN analyses identified only 2262 m6A peaks (FDR cutoff of p<e10^-5^) in *METTL3-KD* hTSCs (**Dataset S2, sheet2**). Furthermore, many of the existing m6A peaks in *METTL3-KD* hTSCs showed low m6A enrichment compared to control hTSCs (**Fig. 6A-B**, **D-E**). This data indicated that m6A modifications on vast majority of transcripts in hTSCs are regulated by METTL3. The m6A peaks on target transcripts in control hTSCs were distributed throughout the genome (**Supp Fig. S9A-B**, **Dataset 3**). However, the gene ontology (GO) cellular processes analyses of m6A-enriched transcripts in control hTSCs overrepresented ribonucleoprotein complex involving RNA-splicing and mRNA-metabolic processes (**Fig. 6F**). HOMER analyses identified that within the m6A peaks, the CUGCAG motif is the most enriched motif in both control (*p*=1e^-209^) and *METTL3KD* (*p*=1e^-70^) hTSCs (**Supp Fig. S9C-D**, **Dataset S4-5**). Analyses of transcripts with differential enrichment of m6A modification in control *vs. METTL3KD* hTSCs identified 2197 transcripts (**Dataset S6**) on which m6A enrichment were either lost or reduced in *METTL3KD* hTSCs. Furthermore, PlacentaCellEnrich analysis revealed that the downregulated m6A peak associated transcripts most significantly represent trophoblast cells of a human placenta (**Fig. 6G**).

**Fig. 6.**
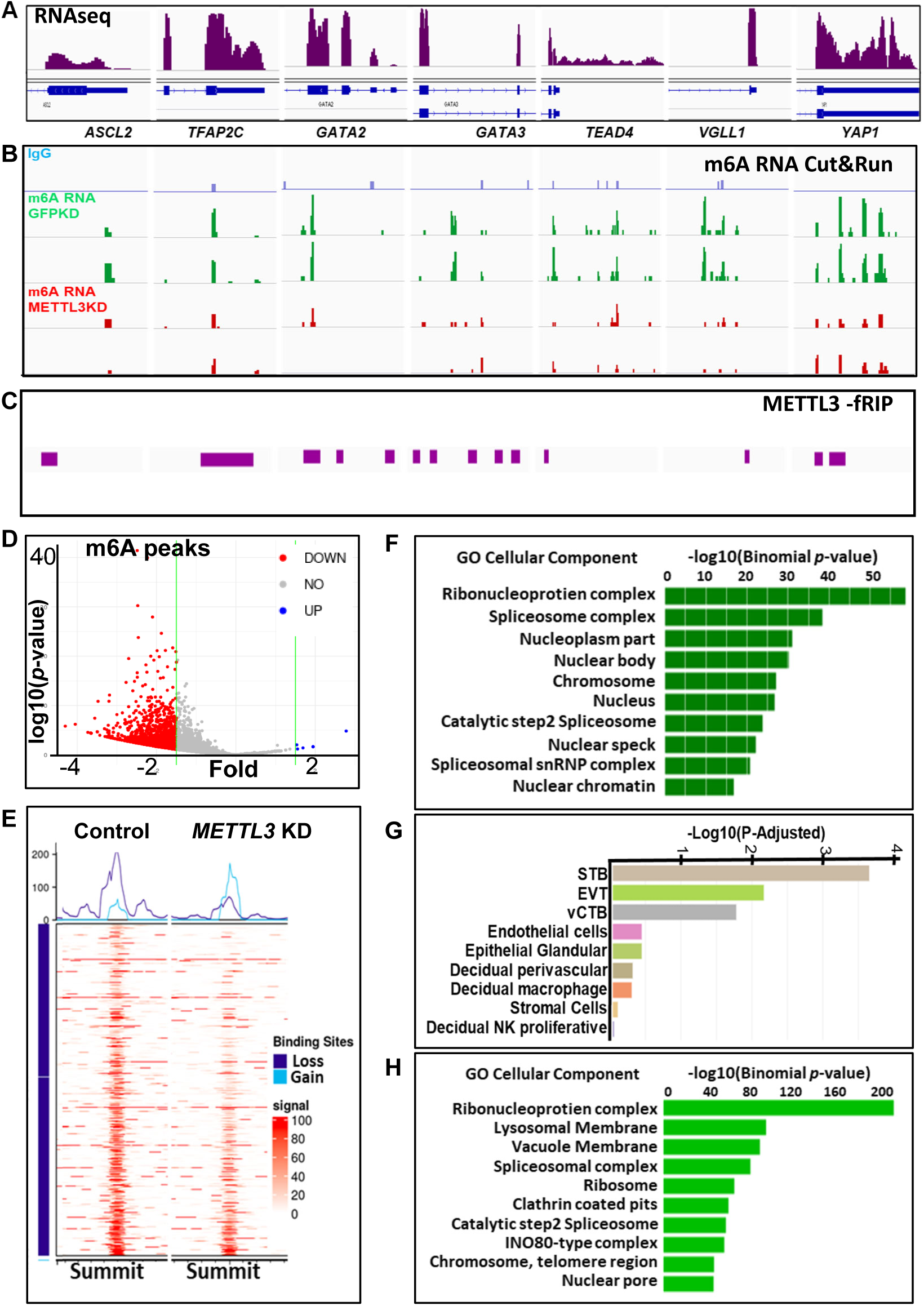
METTL3 shapes m6A RNA modification landscape in hTSC. (A) Integrative Genome Viewer (IGV) tracks showing RNA-seq reads in control hTSC for the *ASCL2*, *TFAP2C*, *GATA2*, *GATA3*, *TEAD4*, *VGLL1* and *YAP1* loci (exon reads are indicated in brown). (B) IGV tracks showing enrichment of m6A modified transcripts (control hTSC in green and *METTL3* KD hTSC in red) for the corresponding genes listed in panel A. It is important to note that several m6A peaks present in control hTSC are either completely lost or significantly reduced in *METTL3* KD hTSC. (C) IGV heatmap showing significantly enriched METTL3 bound regions within the transcripts in the control hTSC (for the corresponding genes in panel A, the heatmap is the average of three independent experiments). (D) Volcano plot showing differentially enriched m6A peaks between controls and *METTL3KD* hTSCs (blue: up-regulated, red: downregulated). (E) Heat maps of differentially enriched m6A peaks (summit ±800bp) between controls and *METTL3KD* hTSCs. (F) GREAT analysis of differentially enriched m6A peaks associated gene showing involvement in regulation of RNA metabolism. (G) PlacentaCellEnrich analysis showing that genes associated with differentially enriched m6A peaks most significantly represent STB, EVT and vCTBs. (H) GREAT analysis of genes corresponding to differentially enriched METTL3-fRIP peaks showing involvement in regulation of RNA metabolism and various nuclear ultrastructure.

To investigate whether m6A-modified transcripts that are differentially regulated in *METTL3KD* hTSCs are direct targets of METTL3, we performed METTL3-fRIP ^57,58^ in control hTSCs. We identified >15000 significantly enriched peaks (FDR cutoff of p<e10^-5^) in comparison to that of IgG negative control (**Fig. 6C**, **Dataset S7, sheet1**) and GREAT tool identified 8615 METTL3 bound transcripts in hTSCs (**Dataset S7, sheet2**). HOMER analyses identified GCAGCUG as the most enriched (*p*=1e^-1322^) METTL3 bound motif (**Supp Fig. S9E, Dataset S8**) in human TSCs. A comparison of m6A enriched peaks and METTL3 target transcripts identified 1694 genes, which are both m6A modified and are direct targets of METTL3 in hTSCs **(Supp Fig. S9F, Dataset S7, sheet3)**. Among these common METTL3 target and m6A enriched transcripts, 271 transcripts were downregulated, and 256 transcripts were upregulated in *METTL3KD* hTSCs **(Supp Fig. S9G, Dataset S7, sheet4-5)**. Functional analyses of downregulated transcripts indicated that they are most significantly associated with RNA splicing and mitochondrial regulation. In contrast, upregulated transcripts were indicated to be involved in transcriptional regulation.

Recent CRISPR screening studies have identified key essential genes within hTSCs ^23,51^ and transcription factor modules that are associated with CTB self-renewal and EVT/STB differentiation ^59^. Our comparative analyses showed that 61 among 619 GRGs, identified by Dong *et al.,* ^51^ are METTL3 targets and m6A modification on those transcripts are either lost or downregulated in *METTL3KD* hTSCs (**Supp Fig. S10A**). We also found that 46 of the 221 hTSC-specific regulators, identified by Shimizu *et al.,* ^23^ (**Supp Fig. S10A**) and 78 of the 256 CTB-specific genes, identified by Chen *et al.,* ^59^ and are also targets of METTL3 (**Supp Fig. S10B**). We also noticed that 28 out of 127 genes in STB regulators and 11 out of 76 genes in EVT regulators are also METTL3 target and enriched in m6A modification in hTSCs (**Supp Fig. S10B**, **Dataset S9)**.

Pathological pregnancies including preterm birth, PE, IUGR/FGR and RPL are often associated with altered gene expression in the placenta. Therefore, we focused on examining the association of METTL3-regulated, m6A modified transcripts in hTSCs along with human pathological pregnancies (**Supp Fig. S10C-F**). Notably, transcripts of 44 out of 429 genes in early preterm placentae ^60^, 34 out of 252 genes in PE placentae ^61^, 73 out of 621 genes in PE-FGR placentae ^62^, and 99 out of 634 genes in recurrent miscarriage (RPL) placentae ^63^ were identified as targets of METTL3 in hTSCs **(Supp Fig. S8F, Dataset S9)**. In conclusion, our comprehensive examination of METTL3-mediated m6A modification in hTSCs provides compelling evidence that METTL3 function is pivotal for the maintenance, proliferation, and differentiation of human trophoblast progenitors through m6A modification and defective METTL3 function could alter gene expression program in developing trophoblast cells leading to pathological human pregnancies.

Importantly, we also noticed that METTL3 target mRNAs, on which m6A modification was reduced in *METTL3KD* hTSCs, include transcripts like *TEAD4*, *YAP1*, *ASCL2*, *GATA2*, *GATA3*, and *TFAP2C*, which are either essential for the maintenance of hTSC self-renewal or EVT development (**Fig. 6A-B**). However, RNA-seq analyses did not show significant downregulation of these genes in *METTL3KD* hTSCs. We reasoned that the RNA-seq data might not be sensitive enough to capture subtle changes in transcript levels of these essential trophoblast regulators. Therefore, we performed RT-qPCR and noticed that transcript levels of all these genes were downregulated by >50% in *METTL3KD* hTSCs (**Fig. 7A**). In addition to these key regulators, RT-qPCR analyses also revealed loss of transcripts for *TP63*, which is essential for hTSC/CTB self-renewal ^64,65^. In contrast, RT-qPCR analyses did not show significant alterations in transcript levels of STB-specific genes *CGB* and *SDC1*. Thus, unbiased analyses of m6A enrichment, METTL3-fRIP along with RT-qPCR analyses indicated that METTL3-mediated m6A modification is essential to maintain transcript levels of key regulators, essential for human trophoblast development and placentation.

**Fig. 7.**
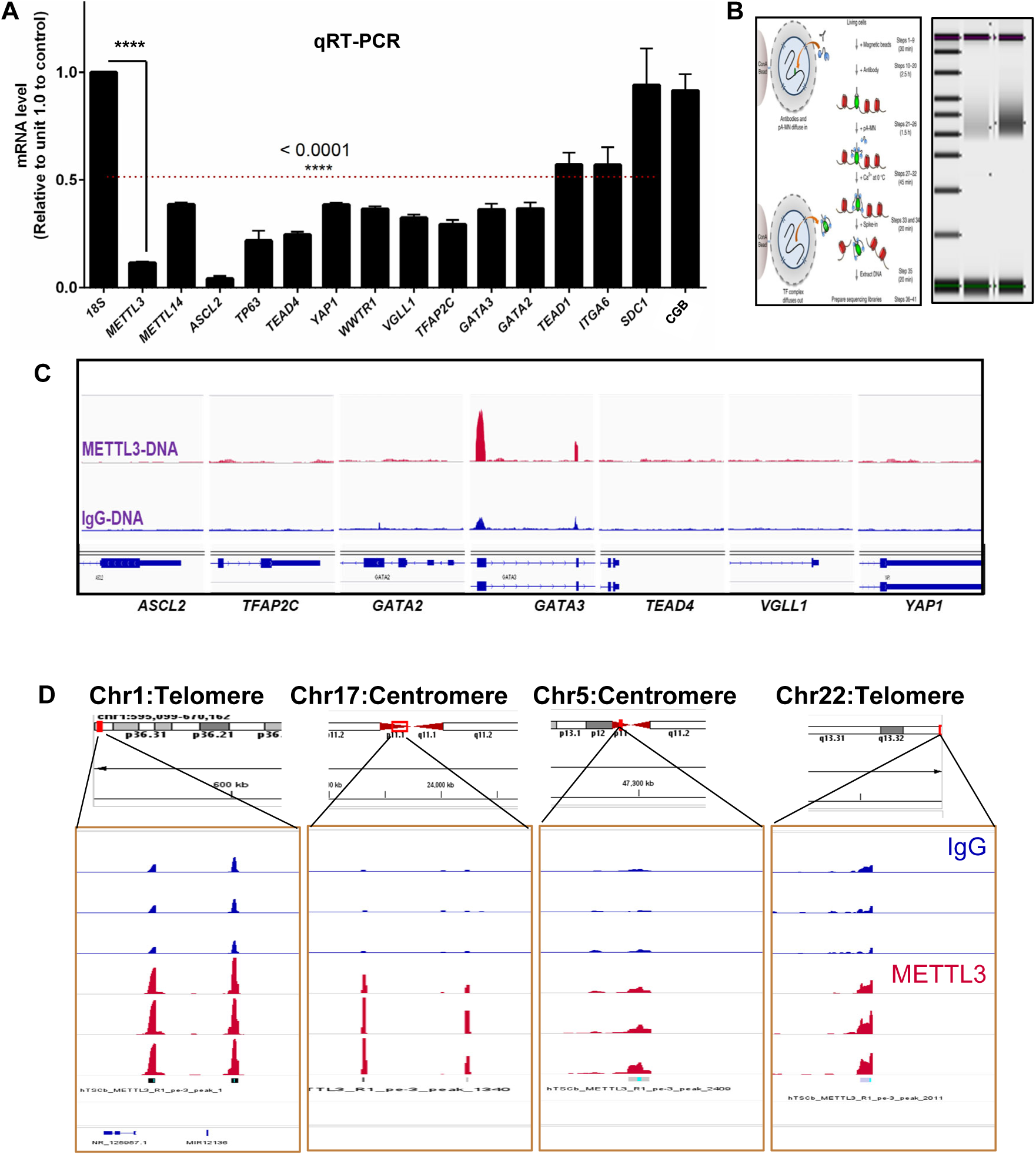
METTL3 DNA binding activity does not regulate transcription of key trophoblast genes. (A) The qRT-PCR plot shows that depletion of METTL3 in hTSCs lead to loss of transcript levels of key trophoblast regulators, on which m6A modification was detected and reduced upon METTL3 depletion. The transcript level for each gene in control TSCs were considered as 1 (n=3 independent experiment, the dotted bar represents gene expression changes with a statistical significance of p<0.0001). Please note that the transcripts for housekeeping gene 18s rRNA as well as STB markers *SDC1* and *CGB* were not significantly altered upon METTL3 depletion. (B) Workflow of the METTL3-CUT&RUN-seq. (C) IGV tracks showing METTL3 binding at the genomic loci of key trophoblast regulators, for which transcript levels were reduced in *METTL3*-KD hTSCs. It is important to note that other than *GATA3*, none of the genomic locus was enriched for METTL3 DNA binding. (D) IGV tracks showing significantly enriched METTL3 bound chromatin regions in hTSCs. Note that the majority of significantly enriched METTL3 peaks are localized at telomere or centromeric regions on the chromosome.

As METTL3 is also known to regulate transcription *via* direct binding to the chromatin, we wanted to understand whether any of the key trophoblast regulators are directly regulated by METTL3 binding at their chromatin domain in hTSCs. Therefore, to uncouple the epigenetic ^56,66^ function of METTL3 from epitranscriptomic m6A modification, we performed CUT&RUN (**Fig. 7B**) ^67,68^ with METTL3-antibody to identify chromatin regions that are direct targets of METTL3 in hTSCs. We identified 350 METTL3 peaks with a *p* value cutoff *p* < 1e^−5^ (**Dataset S10, sheet1**). Among the key trophoblast genes, only *GATA3* gene was a direct target for METTL3 in hTSCs (**Fig. 7C**). We found that the majority of METTL3 DNA bound peaks overlap with centromeres and telomeres regions of the human genome (**Fig. 7D** and **Dataset S10, sheet2-3**). These data indicated that METTL3-mediated regulation of key trophoblast transcripts is dependent on m6A modification and independent of METTL3-DNA interaction. However, given that METTL3 directly binds to chromatin region associated with centromere and telomere on different chromosomes, it is possible that METTL3 DNA binding activity is important to maintain genomic stability in developing human trophoblast progenitors.

### METTL3 is required for the self-renewing trophoblast progenitors during early post-implantation development in mouse

METTL3 expression is conserved from mouse to human trophoblast progenitors (**Fig. 1**) and METTL3 loss in human trophoblast progenitors is associated with a subset of idiopathic RPL. Therefore, we posited that METTL3-mediated regulation of trophoblast progenitor self-renewal is a conserved event in mammals and is a necessary mechanism for early stages of placentation. We tested this by evaluating the importance of METTL3 function in primary TSPCs of an early post-implantation mouse embryo. To define importance of METTL3 function in mouse primary TSPCs, we performed loss-of-function studies with a *Mettl3*-conditional knockout mouse model ^69^. We used a mouse model (*Mettl3^fl/fl^:Ubc^-^Cre^ERT^*^2^) in which *Mettl3* could be conditionally deleted with synthetic estrogen receptor ligand, 4-hydroxytamoxifen (4-OHT). We crossed *Mettl3^fl/fl^; Ubc^-^Cre^ERT^*^2^ male with *Mettl3^fl/fl^* females to confine Cre-expression within the developing conceptus. In a post-implantation mouse conceptus, the self-renewing TSPCs reside within the E5.5-E7.5 placenta primordium, consisting of extraembryonic ectoderm (ExE)/ectoplacental cone (EPC) regions. Therefore, we isolated placenta primordia from ∼E7.5 conceptuses, cultured them *ex-vivo* in FGF4/heparin-containing mTSC culture condition and induced CRE-mediated deletion of *Mettl3* with 4-OHT. We found that loss of *Mettl3* in placenta primordia severely affected expansion of primary TSPCs confirming that METTL3 function is necessary for proliferation/self-renewal of primary TSPCs of a post-implantation mouse embryo (**Fig. 8A-F**).

**Fig. 8.**
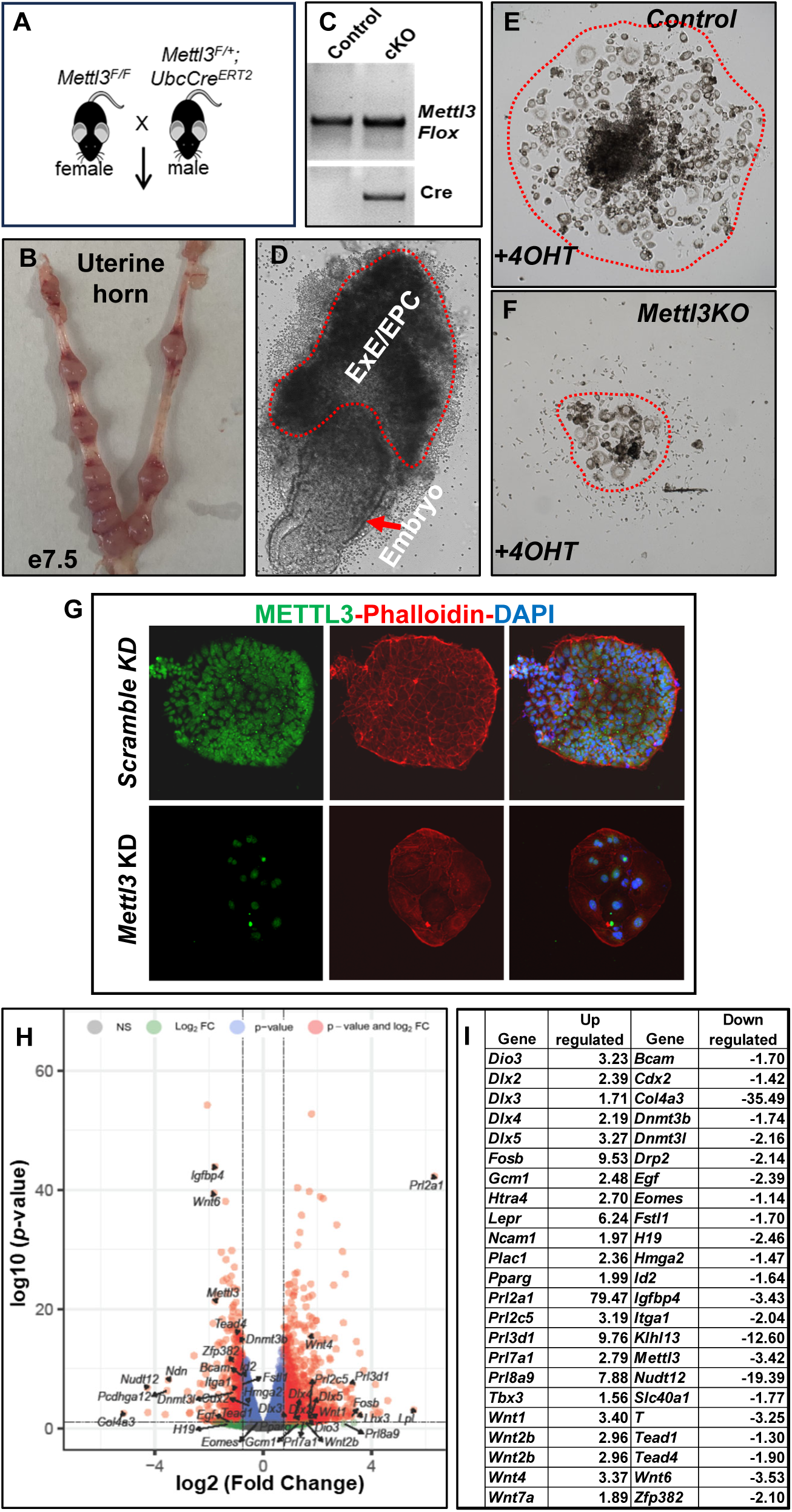
Loss of METTL3 in mouse trophoblast progenitors results in impaired self-renewal potentials. (A) Mating strategy to assay METTL3 requirement in mouse trophoblast progenitors. (B) Representative uterine horn images with ∼E7.5 conceptuses, used for isolation of ExE/EPC regions. (C) Genotyping to confirm expression of *Ubc-Cre^ERT^*^2^ driver. (D) Representative image of ∼E7.5 conceptus, from which the ExE/EPC region were isolated for *ex-vivo* culture. (E) and (F) Representative images of *Mettl3*-KO ExE/EPC cultures, respectively. ExE/EPCs were cultured for 72h upon treatment with 4-hydroxy tamoxifen (4OHT) and FGF4/heparin-containing mouse TSC culture medium. It is important to note that expansion of primary placental progenitors was reduced upon loss of METTL3. (G) Representative micrograph showing loss of METTL3 protein expression in mouse TSCs upon shRNA mediated knockdown of *Mettl3.* The reduction in colony size of *Mettl3*-KD mouse TSCs is also evident. [(METTL3 (green), phalloidin (red) and nuclei were counter stained with DAPI)]. (H) Volcano plot showing DEGs (fold change >1.5 and *p* value < 0.05) between control and *Mettl3KD* mouse TSC. DEGs with important roles in trophoblast development are indicated. (I) Table shows a list of METTL3 target genes, which are important for maintenance of mouse TSC either in proliferative or differentiating state. It is important to note that expression of genes those are required at the proliferative state are downregulated while the expression of genes those are required to initiate differentiation or at the differentiative state are upregulated.

To understand how METTL3 could regulate gene expression program in mouse trophoblast progenitors, we depleted *Mettl3* by RNAi in mouse TSCs (*Mettl3*KD mTSCs) (**Fig. 8G**). Like primary TSPCs of a mouse placenta primordium, the *Mettl3*KD mTSCs showed strong reduction in cell proliferation compared to the control mTSCs (**Fig. 8G**). Furthermore, RNA-seq analyses in control *vs*. *Mettl3*KD mTSCs identified 3348 DEGs (1044 down and 1694 up, log2FoldChange >0.5,) (**Fig. 8H**, **Dataset S11**). Importantly, expression of several mouse TSC stem-state genes, such as *Bcam, Cdx2, H19, Eomes/Tbr2, Tead4, Tead1, Id2, Satb1, Tet1, Sox2, Satb2, Itga1, Zfp382, Hmga2*, *Igfbp4*, *Pvt1*, *Fstl1*, *Wnt6*, *Dnmt3b, Dnmt3l* were down regulated in *Mettl3*KD mTSCs. In contrast, mTSC differentiation markers such as *Gcm1*, *Wnt4, Wnt2b, Wnt1, Wnt2b, Prl3d1/Pl1, Prl8a9, Prl7a1, Prl3d3, Prl3d1* were upregulated in *Mettl3*KD mTSCs (**Fig. 8I**). These observations indicated that, like human TSCs, METTL3 function is essential to balance transcript levels in mouse TSCs to promote self-renewal program and to prevent premature differentiation.

## Discussion

In this study we provide evidence that METTL3, a major m6A RNA methyltransferase ^35–37,70,71^, is essential in two distinct stages of human trophoblast development. First, METTL3 ensures optimum transcript levels for the maintenance of self-renewal in CTB progenitors. Second, METTL3 is essential for EVT development. Using mouse model, we show that *Mettl3-*mediated regulation of trophoblast progenitor self-renewal is a conserved event. Furthermore, our studies with patient-derived hTSCs indicated that a subset of RPL is associated with loss of METTL3 in human trophoblast progenitors. *Mettl3-*mutant mice die ∼E7.5, a developmental stage equivalent to first trimester of placenta. Thus, given the phenotype of METTL3-mutant mice along with our findings that certain idiopathic RPLs are associated with loss of METTL3 in CTB progenitors, it is attractive to propose that impairment of METTL3 expression/function in CTB progenitors of a developing human placenta is a molecular cause for early pregnancy loss.

Over the last few years a number of molecular regulators have been implicated in human trophoblast development ^23,44,51,72–75^. Our mechanistic analyses showed that METTL3-mediated m6A modification is important to maintain optimum transcript levels of several such regulators in hTSCs. We noticed that transcript levels of these regulators either downregulated or upregulated in *METTL3KD* hTSCs. Importantly, mRNA levels of hippo signaling components, transcription factor TEAD4 and cofactors YAP1 and WWTR1, which are essential for hTSC/CTB self-renewal ^20,21,30^, were down regulated in *METTL3KD* hTSCs. METTL3-mediated m6A modification promotes RNA degradation *via* m6A reader proteins, such as YTHDF1-3 and YTHDC1 ^76^. In contrast, m6A modification protects RNA from degradation *via* insulin-like growth factor 2 mRNA-binding proteins (IGF2BPs) ^76^. Gene expression analyses data in human first-trimester trophoblast cells showed that these m6A readers are highly expressed in CTBs and EVTs ^44^. Thus, our data indicate that METTL3-mediated m6A modification acts as an attenuator for fine-tuning the stoichiometry of key transcripts to optimize the hTSC/CTB self-renewal and it would be interesting to define the role m6A-IGF2BP axis in regulation of hippo signaling components during human placentation.

Our loss of function study in hTSCs indicated that METTL3 dictates differentiation fate in hTSCs. Loss of METTL3 in hTSCs induced STB differentiation, indicating that METTL3 functions as gatekeeper to prevent premature STB differentiation in CTB progenitors. Single-cell genomics from our and other laboratories indicated that in a developing human placenta distinct CTB progenitors exist, which can be identified based on their gene expression pattern and replication status ^20,21,77,78^. Given that METTL3 is suppressed in STBs, we propose that METTL3-mediated regulation of transcripts is critical for dictating CTB differentiation landscape and CTB to STB transition during normal placental development. Unlike STBs, METTL3 is essential for EVT development. EVT development is also a multi-stage process, which involves proliferation of column CTBs and presence of intermediate EVT precursors before their maturation to interstitial and endovascular EVTs ^44,79,80^. Our results suggest a bimodal function of METTL3 in human trophoblast progenitors, influencing complex regulatory network that governs gene expression dynamics and cellular fate decisions (**Fig. 9**). Thus, it will be interesting to define what stage/s of EVT development critically relies on METTL3 and underlying mechanism that are dependent on METTL3 function. Analyses of METTL3 targets revealed that METTL3-mediated m6A deposition could be a major regulatory step in RNA metabolism and RNA-splicing in hTSCs. Intriguingly, defective EVT development as well as alternatively spliced gene products such as, soluble FLT1, soluble Endoglin, have been implicated in pregnancy associated disorder, preeclampsia ^81,82^. Furthermore, METTL3 mRNA expression was significantly upregulated in placentae associated IUGR and PE. Thus, it is possible that METTL3-dependent regulation of mRNA-splicing is a key regulatory step during EVT maturation and normal pregnancy outcome.

**Fig. 9.**
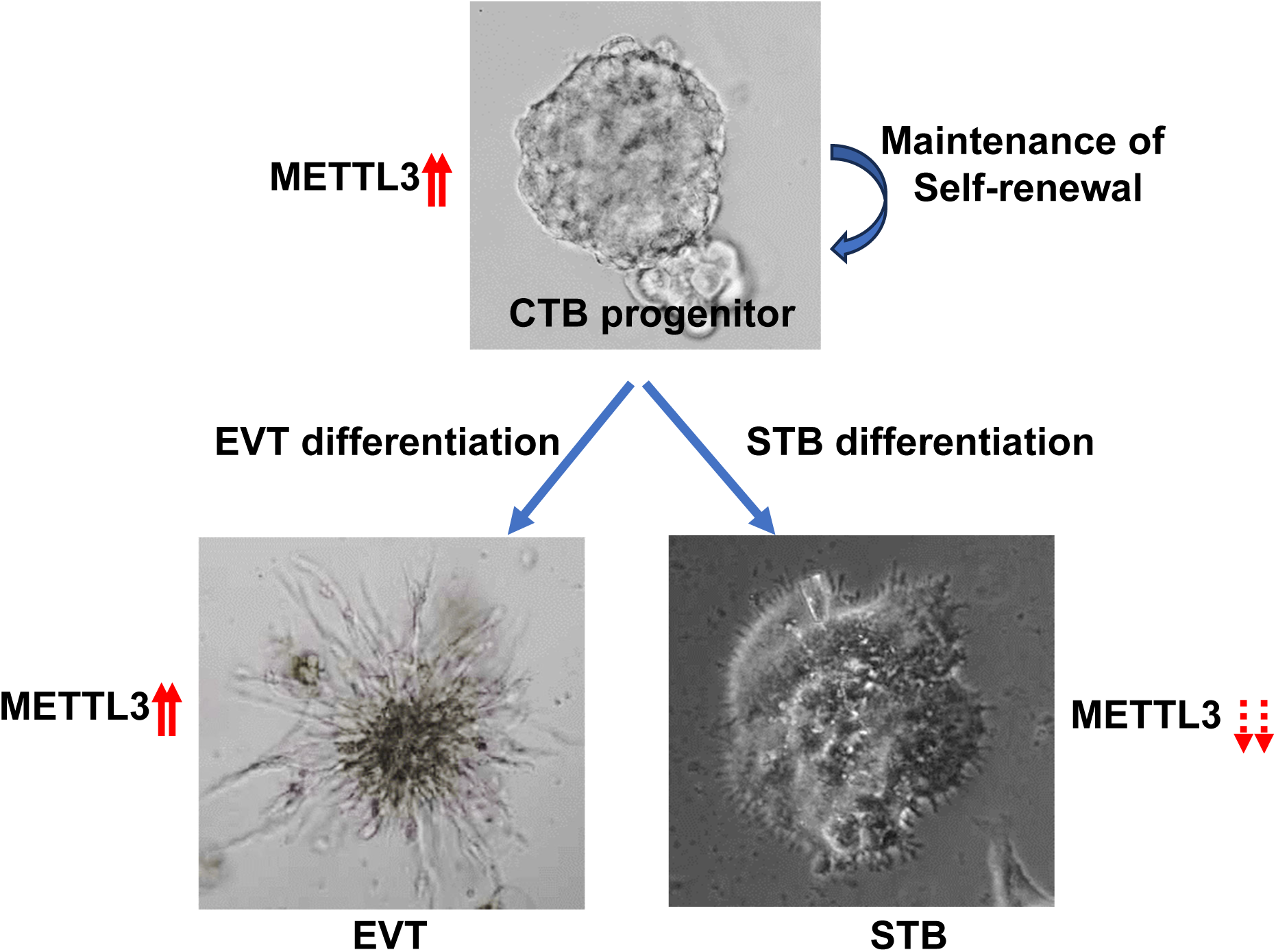
METTL3 expression level governs trophoblast cell fate and lineage differentiation. The model shows that the METTL3 is a key factor in determining the self-renewal ability and differentiation outcome of human CTB progenitors. METTL3 expression is important for the self-renewal of CTB progenitors and their differentiation to the EVT lineage, whereas suppression of METTL3 is important for STB development.

Depletion of METTL3 in hTSCs resulted in loss of m6A enrichment on a vast majority of transcripts, indicating that METTL3 shapes the m6A-epitranscriptomic landscape during human trophoblast development. However, we noticed that m6A modification on several transcripts was not altered in *METTL3KD hTSCs*. It is possible that the residual amount of METTL3 activity in *METTL3KD* hTSCs is sufficient to maintain m6A enrichment on those targets. As hTSCs express other methyltransferases, such as METTL5, METTL16, METTL4, ZCCHC4 and PCIF1 ^83–85^, it is also possible that transcripts that maintain m6A enrichment upon METTL3-depletion are methylated by other methyltransferases. We also noticed that METTL3 was bound to several non-m6A-enriched transcripts, which were differentially expressed in *METTL3 KD* cells, indicating that its role in trophoblast development may be more complex beyond m6A RNA modification. Furthermore, our CUT&RUN analyses indicated that METTL3 binds to numerous chromatin regions, including centromeres and telomeres, in hTSCs. The DNA binding activity of METTL3 might mediate an alternative function, such as maintenance of genomic integrity in human trophoblast cells. Thus, m6A-independent functions of METTL3 might also critically contribute to trophoblast biology during human placentation and further studies are needed to comprehensively elucidate these mechanisms.

## Materials and Methods

### Human TSC culture

hTSC lines, derived from first trimester CTBs following published protocol ^44^ or available hTSC female cell line CT27 ^44^, were cultures exactly as described earlier for maintaining stem state as well as differentiation state of hTSC ^20,21,44^.

### Establishment of hTSC from Recurrent Pregnancy loss (RPL)

RPL placental samples were washed in phosphate buffer saline (PBS) followed by DMEM/F12(Gibco) media and divided into smaller pieces under the dissection microscope in sterile conditions and cultured as described previously ^20,47^. These RPL hTSC were cultures exactly as described earlier for maintaining stem state of hTSC ^20,21,44^ for 3-4 passages until the residual fibroblasts were lost. The established RPL hTSC were cultured together to assess the growth dynamics compared to that of hTSC CT27 ^44^. These cells were also cultured on collagen IV coated coverslips for immunostaining.

### Placental explants culture

First-trimester placental explants were washed in phosphate buffer saline (PBS) followed by DMEM/F12(Gibco) media and divided into smaller pieces under the dissection microscope in sterile conditions and cultured as described previously ^20,47^. All the explant pieces were encapsulated in the Growth factor reduced Matrigel (Corning) mixed with DMEM/F12 (1:1) and left on ice. The placental explant suspended in matrigel was dropped (30-40μl) on the 24 well tissue culture plate. The plate was then inverted to make hanging drop and incubated at 37°C for 20-25 minutes in a humidified chamber in a 5% CO2 for the matrigel-suspension to solidify and encapsulate the explant. Finally, 300μl of EVT media was added to each well and allowed to placental progenitor cells to differentiate into EVT. EVT media was changed as per protocol ^20,44^. The explants were divided into two groups, one group was incubated with DMSO while the other group was incubated with METTL3 inhibitor STM2457 (final concentration 10μM, https://www.selleckchem.com) from day2 onwards. These explants were cultured on for 6-10 days.

### Isolation of vCTB from elective normal placentae

Villous cytotrophoblasts (vCTB) were isolated from the first trimester placentae using two consecutive digestion steps as described earlier^20,48^. Precisely, single placental tissue (8.3 week of gestation) was washed with Mg2+/Ca 2+ -free Hanks balanced salt solution (HBSS, Sigma-Aldrich; H4641) and cut into small pieces. During this process, blood clots, maternal decidua as well as other parts of the placenta were removed completely. Placental villi were collected by centrifugation and incubated for two consecutive digestions with 1x HBSS containing 0.125% trypsin (Gibco; 15090–046) and 0.125 mg/mL DNase I (Sigma-Aldrich; DN25) at 37 °C in the incubator. Digestion was stopped using 10% (vol/vol) FBS and cells were filtered through a 70-μm cell strainer (BD Biosciences). Cells were pelleted by gradient centrifugation. Cells were incubated biotinylated-HLA-G antibody for 10 minutes at RT and HLA-G+ EVTs were selected from the cell suspension by a magnetic stand, the cells bound to the Streptavidin-Biotinylated HLA-G magnetic beads were separated from the suspension. The flow through cell suspension was further processed for ITGA6 following the vender’s prescription (STEMCELL # 17664).

ITGA6 +ve vCTB selected cells were washed twice with 500μL of bTOM on the magnetic stand and residual media was removed and vCTB were processed as per requirement.

### Human TSC organoid generation and differentiation

hTSC, *shScramble* hTSC, *shMETTL3* hTSC*, tetOshMETTL3* hTSC and first trimester primary vCTB were used to generate the self-renewing organoids following earlier described protocols^47,49,86^. Human TSCs of required genotype were harvested and re-suspended in ice-cold basic trophoblast organoid medium (b-TOM containing advanced DMEM/F12 supplemented with 10mM HEPES (Sigma H3537), B27 (Gibco17504-044), N2 (Gibco 17502-048) and 2mM glutamine (Gibco 25030081)). Growth factor reduced matrigel (Corning) was added to the b-TOM cell suspension to reach a final concentration of 60-70%. 30-35µl of the viscous cell solution containing 2×10^3^ cells was dropped in the center of a 24-well plate and then turned upside down and kept at 37°C for 20-25 minutes to generate hanging drop. Finally, the plates are returned to their upward position and the domes are overlaid with 500µl of room temperature a-TOM medium (b-TOM supplemented with 100ng/ml R-spondin (PeproTech 120-38-20UG), 1μM A83-01 (Sigma SML0788), 100ng/ml recombinant human epidermal growth factor (rhEGF, Sigma E9644), 50ng/ml recombinant murine hepatocyte growth factor (rmHGF, PeproTech 315-23-20UG), 2.5μM prostaglandin E2 (R&D System 2296/10), 3μM CHIR99021 (Sigma SML1046) and 100ng/ml Noggin (Invitrogen PHC1506)). The organoids are allowed to grow for 8-10 days with fresh media being changed every 3 days supplemented with appropriate drugs (as indicated in result sections). Brightfield images were taken to observe the growth of the organoids. For EVT differentiation, P1 organoid were trypsinized to form single cell suspension, washed with bTOM media and replated 2×10^3^ cells in a 24-well plate as 4-5 (10µl) domes/well and then turned upside down and kept at 37°C for 20 minutes to generate hanging drop and allowed to form very small organoid for 2-3 days. Once the organoid formed, a-TOM media was replaced with EVT media as described earlier^20,44^ and treated with required drugs.

### Single-Cell RNA sequencing and analysis

scRNA-seq analysis with first-trimester placenta was performed and detailed analysis were reported earlier ^20,21^. Single-cell RNA-seq data used to generate METTL3 expression in human TSC and STB were performed and detailed analysis were reported earlier^23^.

### Short hairpin RNA (shRNA) mediated RNA interference

To generate *METTL3* knockdown human TSCs and mouse TSC, lentivirus particles carrying shRNA against *METTL3* mRNA were used. A scramble shRNA with sequence was used as control. The TSC were treated with 8μg/ml polybrene prior to transduction. Cells were selected in the presence of puromycin (1-2µg/ml). Selected cells were tested for knockdown efficiency and used for further analyses. Freshly knocked-down cells were used for each individual experimental set to avoid any silencing of shRNA expression due to DNA-methylation at LTR under continuous puromycin selection. To generate data at least 3-4 individual experiments were done to get statistically significant results.

### *hMETTL3* overexpression

To generate overexpression *METTL3* in human TSCs, lentivirus particles carrying a construct carrying RNA against *METTL3* mRNA (pCDNA-FLAG-METTL3, #160250) was used. A scramble shRNA with sequence was used as control. The TSC were treated with 8μg/ml polybrene prior to transduction. Cells were selected in the presence of puromycin (1-2µg/ml) or Gentamycin (100µg/ml).

### Human placental tissue sample analyses

Formaldehyde fixed, de-identified first trimester as well as other pathological placental tissues were obtained from Mount-Sinai hospital, Toronto. Term Placental tissues were obtained at the University of Kansas Medical Center with consent from patients.

### Cell proliferation assay

Human TSCs were seeded (2,000cells/well of 12 well plate) on collagen IV coated coverslips and cultured for 72 hours to assess cell proliferation. Before harvesting coverslips, cells were treated with BrdU in the cell culture medium for 45 minutes. Cell proliferation was assessed using BrdU labeling assay and detection kit (Roche Ref#11296736001) in live cells following manufacturers’ protocol. After anti-BrdU staining, hTSC colonies were imaged using Nikon 90i and manually counted the total number of DAPI and BrdU positive nuclei per field.

### mRNA expression analyses by RT-PCR

Total RNA was extracted from the cells, human placentae, mouse placentae using RNeasy Mini Kit (Qiagen-74104) using manufacturer’s protocol. cDNA was prepared from total RNA (1μg). Primer cocktail comprising of 0.2μg/μl oligo dT and 50ng/μl random hexamer was annealed to the RNA at 68° for 10 minutes, followed by incubation with the master mix comprising of 5X first strand buffer, 10mM dNTPs, 0.1M DTT, RNase Inhibitor and M-MLV transcriptase (200U/μl) at 42^°^ for 1 hour. The cDNA solution was diluted to 10ng/μl and heat inactivated at 95^°^ for 5 minutes. Real-time PCR was performed using oligonucleotides (listed below). 20ng equivalent of cDNA was used for amplification reaction using Power SYBR Green PCR master mix (Applied Biosystems-4367659).

### RNA-seq analyses

Total RNA was used to construct RNA-seq libraries using the Illumina TruSeq Stranded Total RNA Sample Preparation Kit according to manufacturer’s instructions. RNA seq was performed using Illumina HiSeq 2500 platform. The detailed protocol is mentioned in SI Appendix, Supplementary Materials and Methods.

### m6A RNA CUT&RUN

Total RNA was isolated and purified using RNeasy Mini Kit (Qiagen-74104) using manufacturer’s protocol where the RNA was treated with DNase I on the column. Purified RNA were processed for m6A RNA CUT&RUN ^87^ using EpiNext CUT&RUN RNA m6A-Seq Kit (P-9016-12). In brief, total RNA (5 µg) extracted from *tetOshGFP* and *tetOshMETTL3* hTSC was subjected to immunoprecipitation with the m6A antibody and IgG (P-9016, EpiGentek, 1:100 dilution) and cleaved on beads. The beads were then washed, RNA was purified from the beads and subjected to indexed library preparation following the vendors prescribed protocol. The libraries were sequenced with a NovaSeq 6000 system (Illumina). Additional details of CUT&RUN-seq data analyses are mentioned in SI Appendix, Supplementary Materials and Methods.

### METTL3 CUT&RUN

Proliferating semiconfluent 200,000 live hTSC were used per sample for CUT&RUN following published protocol ^67,68^. Cells were captured on Concanavalin A–coated beads (EpiCypher, Durham, NC); cell permeabilization was done using buffers containing 0.5% wt/vol Digitonin before incubation with anti-METTL3 antibody. Protein A and G fused Micrococcal Nuclease (EpiCypher) was used for DNA digestion. The detailed protocol is mentioned in SI Appendix, Supplementary Materials and Methods.

### Statistical Analyses

Statistical significance was determined for quantitative RT-PCR analyses for mRNA expression and for other quantitation analyses. We performed at least n=3 technical or biological replicates for all these experiments. For statistical significance of generated data, statistical comparisons between three means were determined with Student’s t test, and significantly altered values (*p* ≤ 0.01) are highlighted in the figures by an asterisk. RNA-seq data were generated with n = 2-3 experimental replicates per group. The statistical significance of altered gene expression (absolute fold change ≥ 1.5 and false discovery rate *q*-value ≤ 0.05) was initially confirmed with right-tailed Fisher’s exact test. Independent datasets were analyzed using GraphPad Prism software.

### Ethics Statement regarding studies with mouse model and human placental tissues

All studies with mouse models were approved by IACUC at the University of Kansas Medical Center (KUMC). Human placental tissues (6^th^-9^th^ weeks of gestation) were obtained from legal pregnancy terminations *via* the service of Research Centre for Women’s and Infants’ Health (RCWIH) BioBank at Mount Sinai Hospital, Toronto, Canada. The Institutional Review Boards at the KUMC and at the Mount Sinai hospital approved utilization of human placental tissues and all experimental procedures.

## Author Contributions

Soumen Paul and Avishek Ganguly conceived and designed the initial experiments. Ram Kumar redesigned and performed all the experiments, analyzed data, and wrote manuscript. Ananya Ghosh and Md. Rashedul Islam performed experiments. Rajnish Kumar performed genomic sequence data processing and downstream bioinformatics analyses. Ram Kumar performed all bioinformatics analyses after sequence data processing. Soma Ray, Abhik Saha, Namrata Roy, Taylor Knowles, Purbasa Dasgupta, Asef Jawad Niloy helped in establishing RPL hTSC. Courtney Marsh helped in providing placental samples.

## Acknowledgement

This research was supported by NIH grants HD062546, HD0098880 and HD079363 to SP. RPK was partially supported from the Department of Pathology Internal Medicine internal funds. This study was also supported by various core facilities, including the Genomics Core, Imaging, and histology Core facility of the University of Kansas Medical Center. The *Mettl3* floxed mice was a generous gift from Dr. Jacob H Hanna. We also acknowledge administrative support from the Department of Pathology Internal Medicine and the Institute for Reproduction and Perinatal Research.

## Data availability

All the sequencing data are available and will be released in a public database upon acceptance of this manuscript.

## Competing Interest Statement

Authors do not have any conflict of interest to disclose.

## Notes

### Competing Interest Statement

The authors have declared no competing interest.

