## Supporting Figures for "METTL3 shapes m6A epitranscriptomic landscape for successful human placentation"

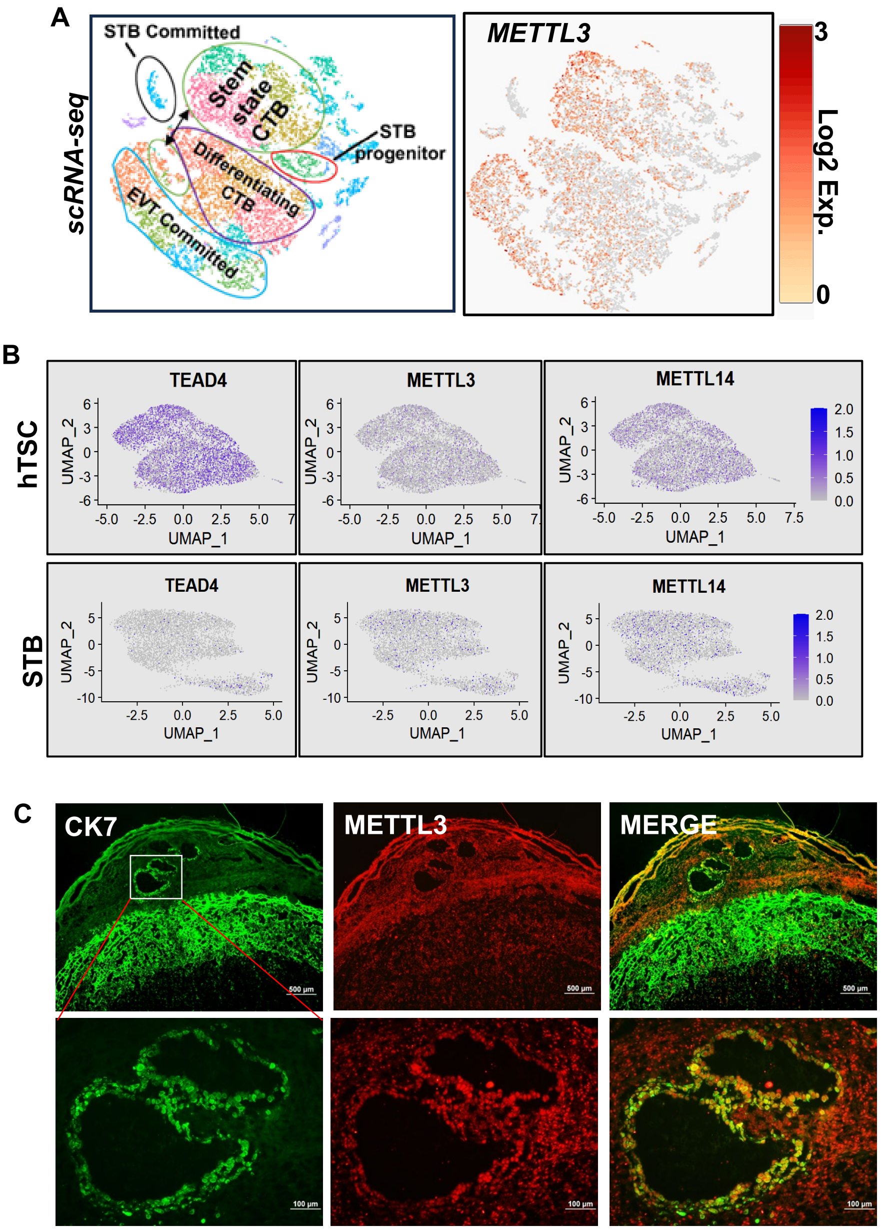


**Fig. S1.** (A) Differential mRNA expression patterns of METTL3 in single-cell clusters, obtained by scRNA-seq analyses in first-trimester human placentae (Data reanalyzed from previous publication, Saha et al., PNAS 2020, Ref. 21 of main manuscript). t-SNE plots showing clusters of stem-state CTBs (green ellipticals), immature STBs (red elliptical), mature STBs of cluster (black elliptical), differentiating CTBs (brown elliptical) and committed EVTs (light blue shape). (B) scRNAseq analysis showing METTL3 expression in hTSC (upper panel) and differentiated STB (lower panel) along with TEAD4 and METTL14. Data reanalyzed from previous publication (Wang *et al.* Nature Genetics 2024, Ref. 45 of main manuscript). (C) Immunofluorescence images of histological sections of an E14.5 rat uterine-placental interface showing expressions of METTL3 (red) along with Cytokeratin (CK7, green). The lower panels represent an enlarged section of the uterus showing METTL3 expression in invasive endovascular trophoblast cells, lining of the uterine artery.

**Fig. S2**

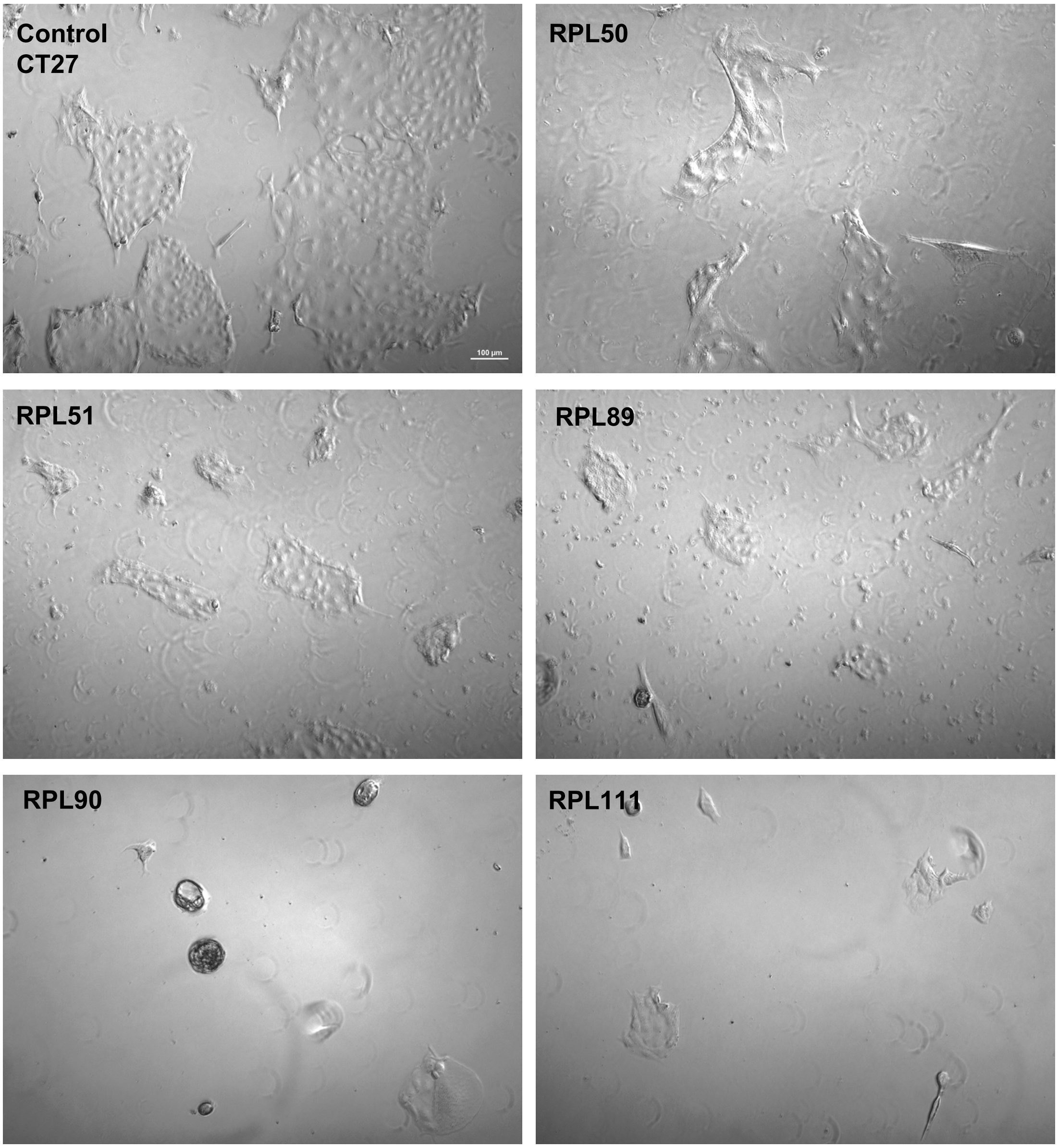


**Fig. S2.** hTSC lines established from human placenta associated with idiopathic recurrent pregnancy loss (RPL). Representative micrographs are shown for CT27 hTSC (control, day 3) and RPL hTSC lines (day 6). 1x104 number of cells were seeded on a 12 well plate and the cell colonies were imaged and compared using a brightfield microscope on different day points (scale 200μm). Images show that unlike the control CT27 hTSCs, the colonies of RPL TSC lines are extremely slow growing.

Fig. S3

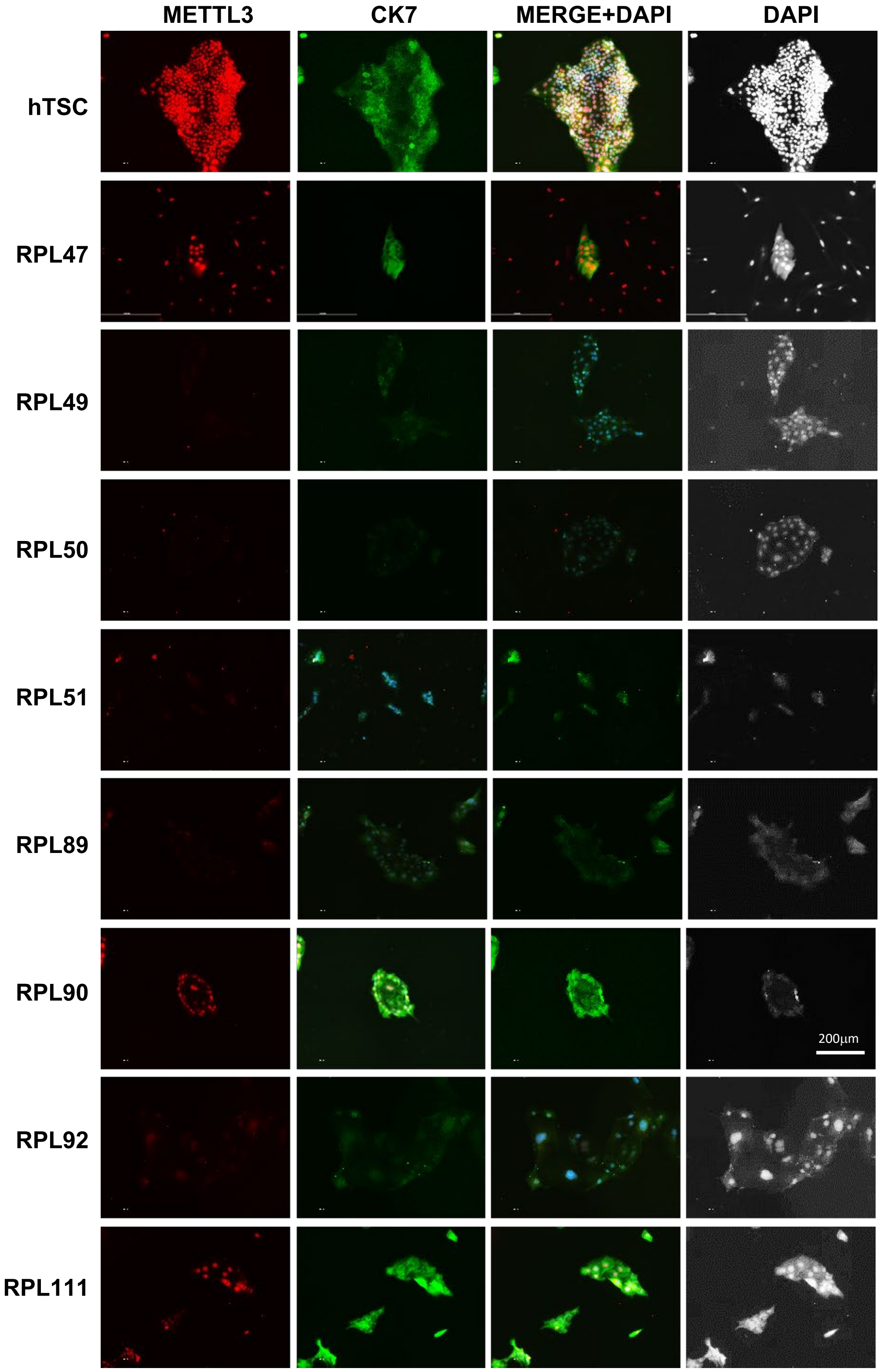


**Fig. S3.** Immunohistochemistry showing expression of METTL3 protein levels in the control CT27 hTSC and established RPL hTSC lines. Note that RPL colonies are extremely slow in proliferation and show very low level of METTL3 protein expression compared to the control CT27 hTSCs.

Fig. S4

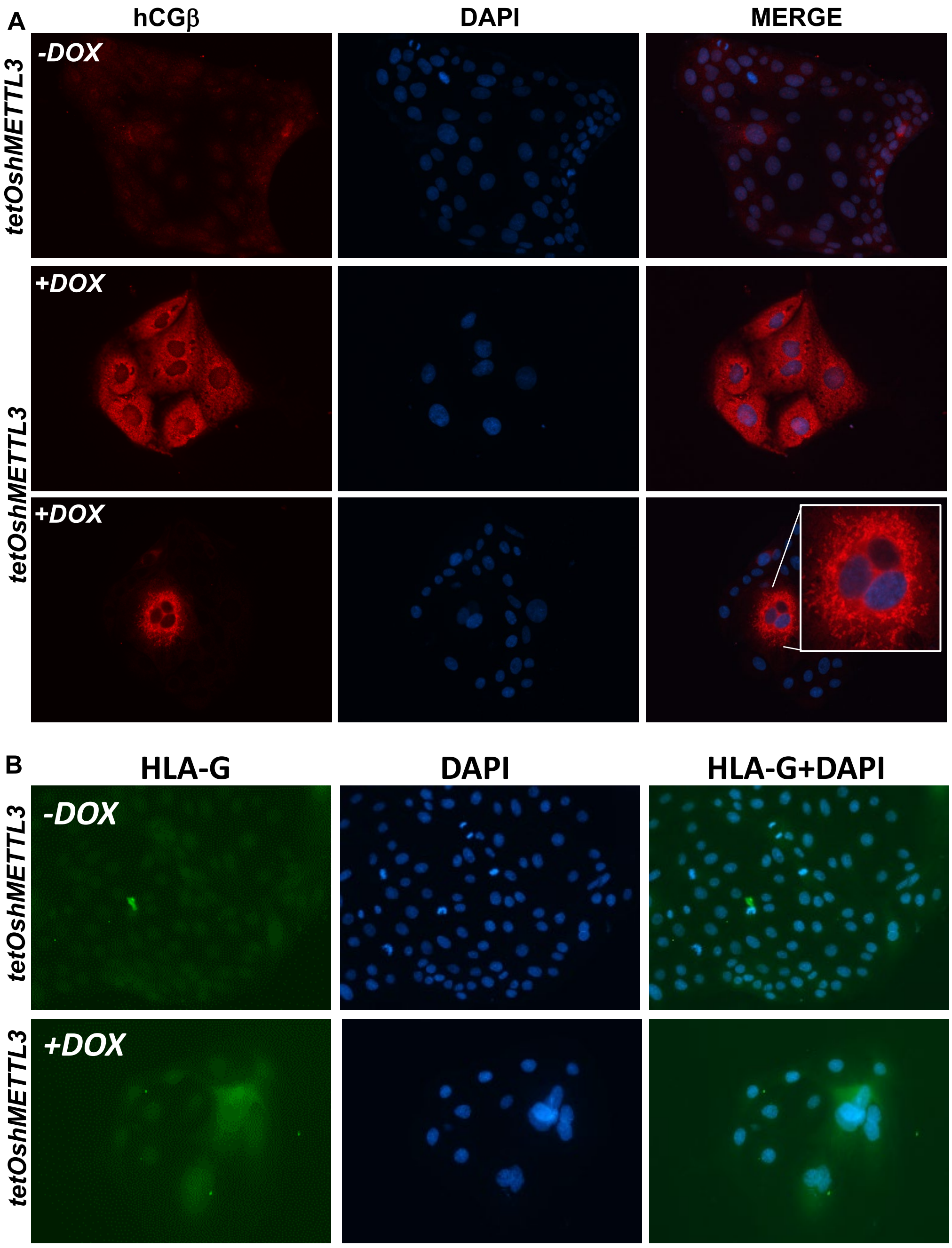


**Fig. S4. METTL3 protects premature STB differentiation in HTSCs.** (A) Immunofluorescence images show that depletion of METTL3 in CT27 hTSCs induces expression of hCGβ, a marker for STB differentiation. Two different knockdown colonies are shown. (B) Immunofluorescence images show that loss of METTL3 does not induce EVT differentiation, evident from the lack of HLA-G expression.

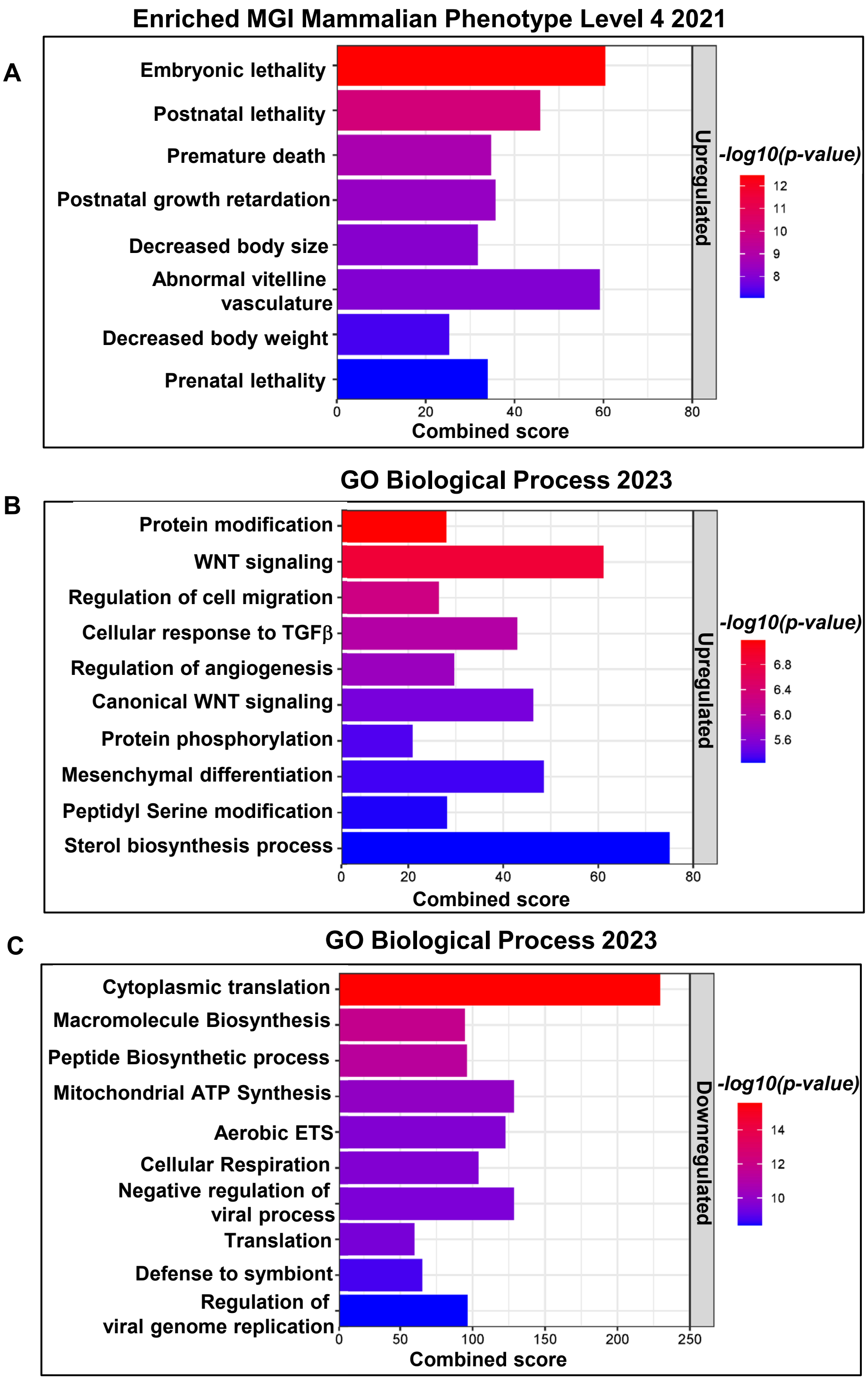


**Fig. S5**. EnrichR analysis of METTL3-regulated differentially expressed gene associated pathways.

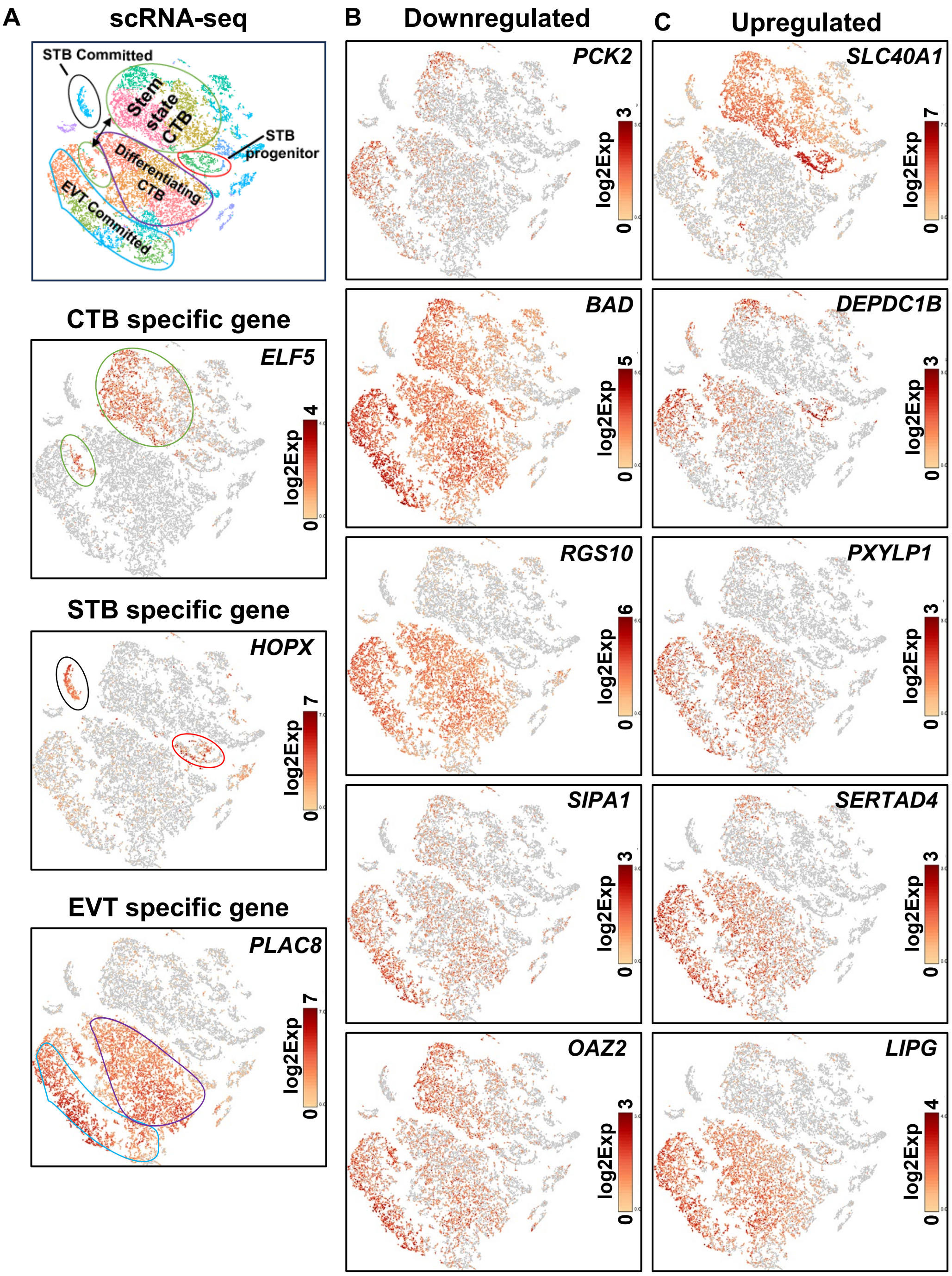


**Fig. S6.** (A) Single cell clusters obtained by scRNA-seq analyses in first-trimester human placenta. Clusters representative of undifferentiated CTBs, and CTBs committed to STB and EVT differentiation are shown (Data reanalyzed from previous publication, Saha et al., PNAS 2020, Ref. 21 of main manuscript). (B-C), Expression patterns of METTL3-regulated key genes in human first-trimester single cell clusters representing CTBs stem-state or differentiation to EVT or STB lineages.

Fig. S7

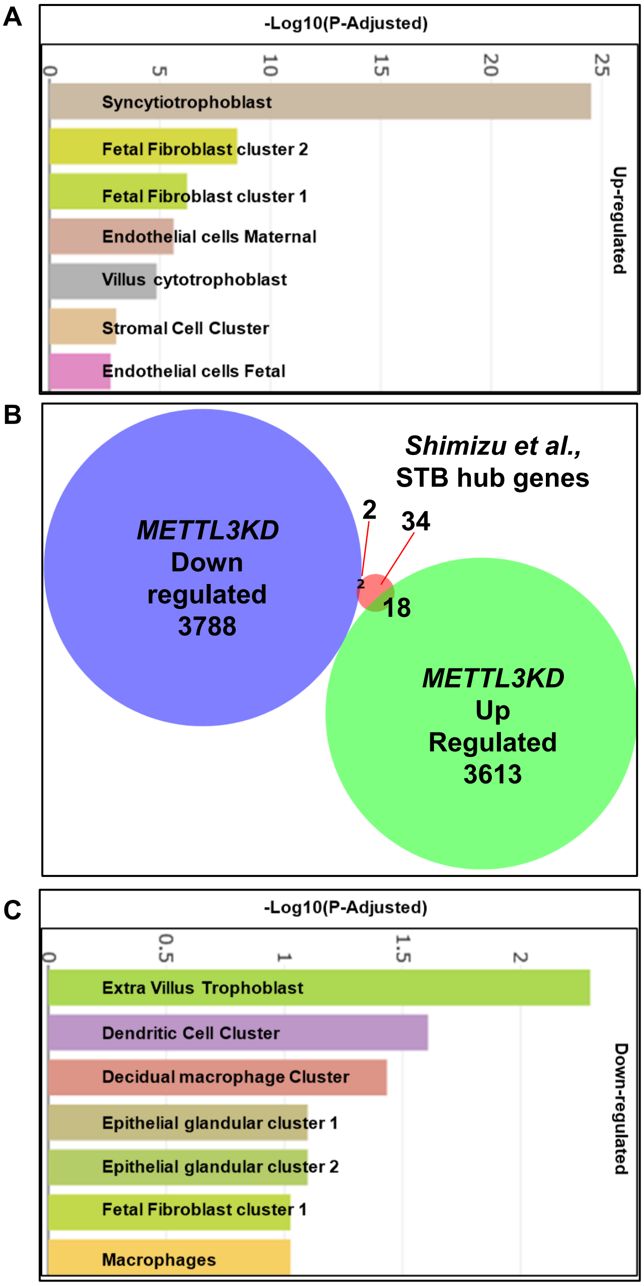


**Fig. S7**. (A) PlacentaCellEnrich analysis of METTL3-regulated differentially expressed gene in METTL3KD hTSCs. It is important to note that the STB-associated genes are most robustly upregulated. (B) Venn diagram showing 18 genes out of 54 STB hub-genes, identified by Shimizu et al., PNAS 2023 (Ref. 23 of main manuscript) were upregulated in METTL3KD hTSCs. (C) PlacentaCellEnrich analysis of down regulated gene showed strong association with EVT development.

Fig. S8

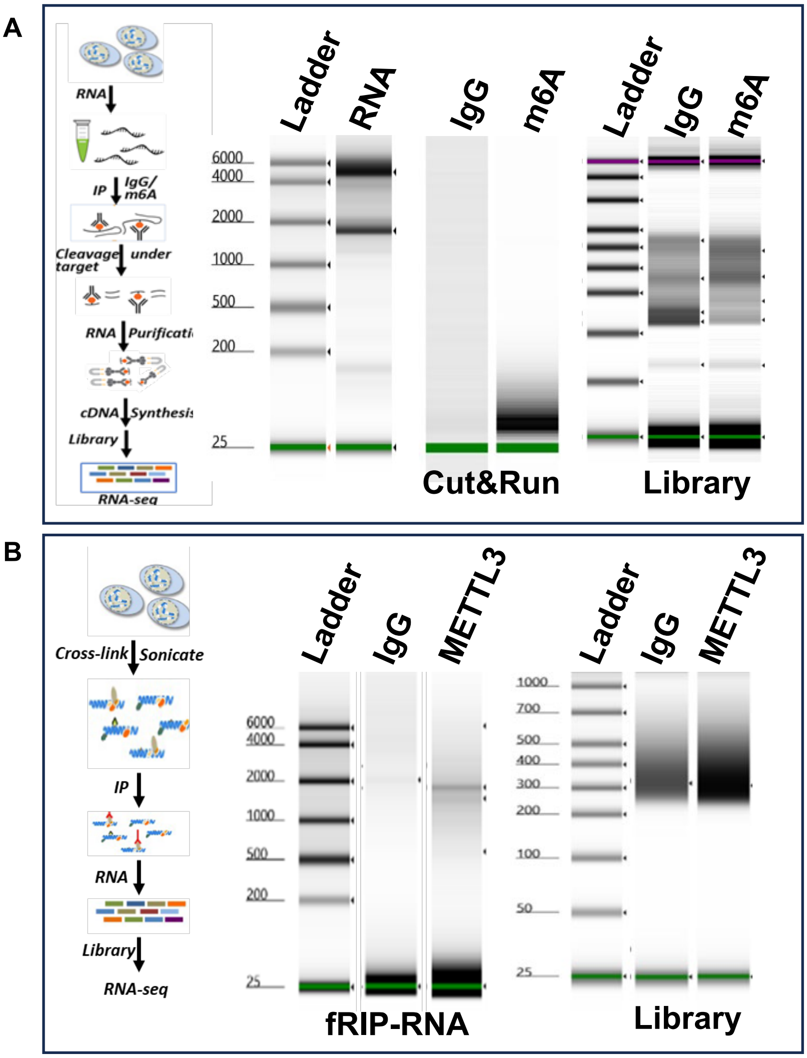


**Fig. S8**. (A) Experimental flow showing m6A-RNA Cut&Run and (B) f-RIP.

**Fig. S9**

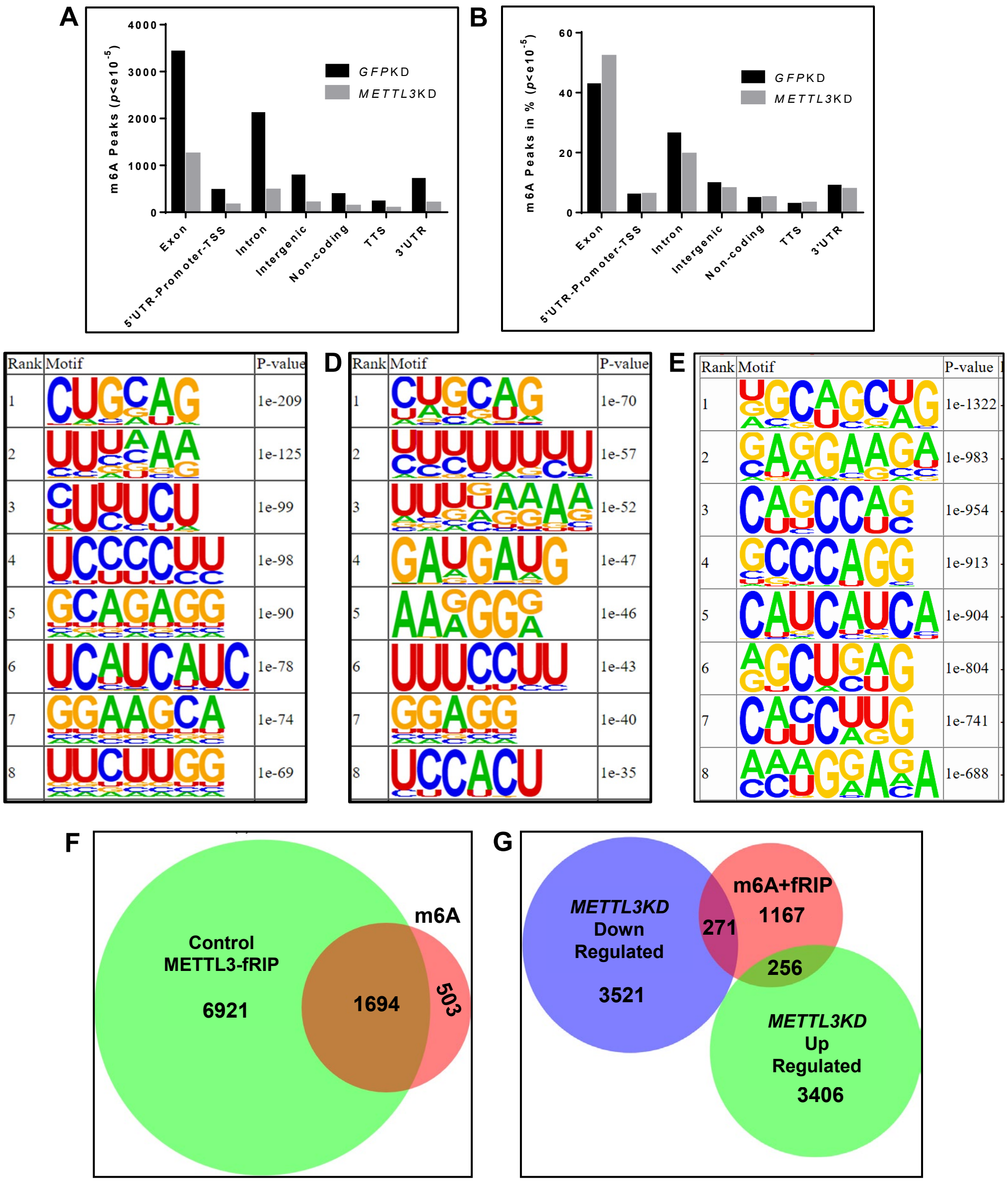


**Fig. S9**. METTL3-mediated regulation of m6A modification on transcripts within hTSCs. (A-B) Bar chart showing the genomic distribution of m6A modified peaks between controls and *METTL3*KD hTSC. (C-D) Homer analysis showing most enriched METTL3 associated RNA binding motifs within the m6A modified regions. (E) Homer analysis showing most enriched METTL3 associated RNA binding motifs within the METTL3-RNA binding regions. (F) Venn diagram showing the number of METTL3 regulated m6A modified transcripts that are significantly altered in *METTL3*KD hTSCs. (G) Venn diagram showing the number of m6A modified transcripts directly bound by METTL3 and are significantly altered in *METTL3*KD hTSC.

Fig. S10

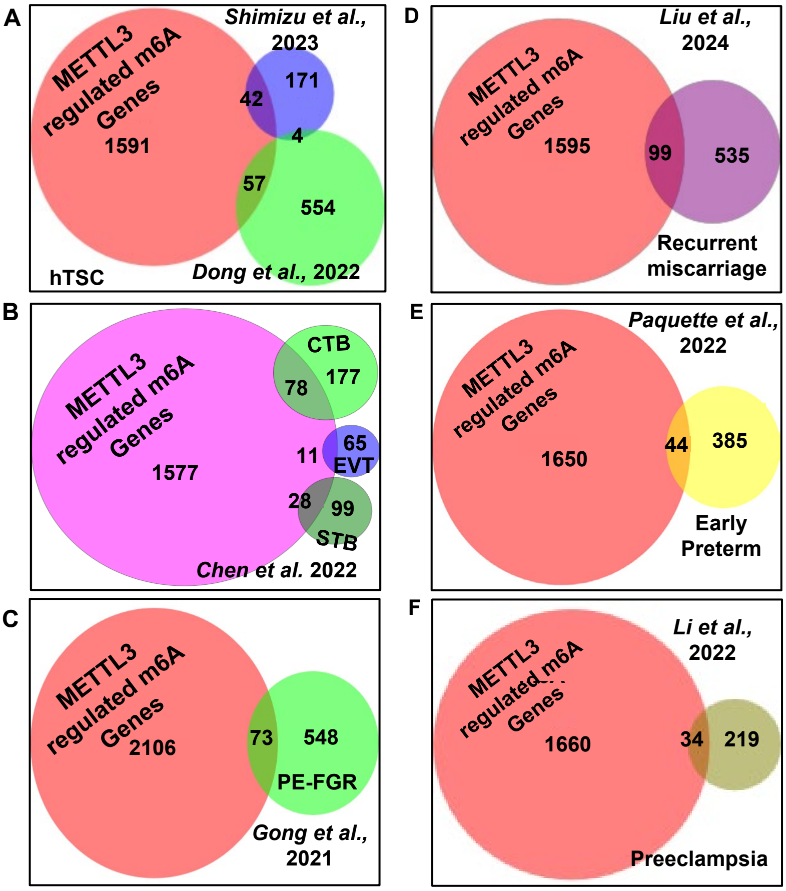


**Fig. S10**. Venn diagram showing the number of METTL3 regulated m6A modification associated transcripts (m6ADiff Genes) and association with human TSC (A), normal placenta (B) and placental pathological conditions (C-F).
