## Supporting DATA for "METTL3 shapes m6A epitranscriptomic landscape for successful human placentation"

**Material and methods:**

**List of primary antibodies:**

| **Name** | **Company** | **Catalog** |
|---|---|---|
| METTL3 | Proteintech | #15073-1-AP |
| METTL3 | CST | #86132 |
| m6A | CST | #56593 |
| m6A | Active motif | #61756 |
| Pan-cytokeratin | Abcam | ab9377 |
| β-Actin | Sigma | A5441 |
| HCGβ | Abcam | ab53087 |
| HLA-G | Abcam | ab52455 |
| HLA-G | Abcam | ab239332 |
| Biotin-HLA-G | Abcam | ab52456 |
| OCT4 | Santa Cruz | sc-5279 |
| Cytokeratin 7 | Dako | M7018 |
| E-cadherin | Abcam | ab1416 |
| IgG | CST | # 66362S |
| LaminB | Santa Cruz | sc-6216 |
| H3K4me3 | CST | C42D8 |
| CD49f-PE | STEMCELL | # 17664 |
| Dynabeads | Invitrogen | # 65601 |

**List of secondary antibodies:**

| **Name** | **Company** | **Catalog** |
|---|---|---|
| Alexa fluor 488 goat anti-rabbit IgG | Invitrogen | A11008 |
| Alexa fluor 568 goat anti-mouse IgG | Invitrogen | A11031 |
| Alexa fluor 488 donkey anti-mouse IgG | Invitrogen | A21202 |
| Alexa fluor 568 donkey anti-rabbit IgG | Invitrogen | A10042 |
| Alexa fluor 488 donkey anti-Goat IgG | Invitrogen | A-11055 |

**Genotyping primers:**

| | |
|---|---|
| *Mettl3 flox* | GTTGATGAAATTATCAGTACAATGGTTCTGA |
| | GTAAAGAACAACTCTGGTTATCGTCATCG |
| *Cre* | AAAATTTGCCTGCATTACCG |
| | ATTCTCCCACCGTCAGTACG |

**RT-PCR human primers:**

| **Gene** | **Forward Primer** | **Reverse Primer** |
|---|---|---|
| *HPRT1* | ACCCTTTCCAAATCCTCAGC | GTTATGGCGACCCGCAG |
| *18SRNA* | AAC CCG TTG AAC CCC ATT | CCA TCC AAT CGG TAG TAG CG |

| | | |
|---|---|---|
| *hMETTL3* | CATTGCCCACTGATGCTGTG | AGGCTTTCTACCCCATCTTGA |
| *hMETTL14* | AGTGCCGACAGCATTGGTG | GGAGCAGAGGTATCATAGGAAGC |
| *hCGB* | GTGTGCATCACCGTCAACAC | GGTAGTTGCACACCACCTGA |
| *hTP63* | GTCATTTGATTCGAGTAGAGGGG | CTGGGGTGGCTCATAAGG T |
| *hTEAD4* | ACGGCCTTCCACAGTAGCAT | CTTGCCAAAACCCTGAGACT |
| *hVGLL1* | TCAGAGTGAAGGTGTGATGCT | GCACGG TTTGTGACAGGTACT |
| *hITGA6* | GGCGGTGTTATGTCCTGAGTC | AATCGCCCATCACAAAAGCTC |
| *hTFAP2C* | TACTGGGAGGTGTTCTCAGAAG | GGCCGGAAGATTCAACCCAAT |
| *hASCL2* | AACTTGAGCTGCTGGAGGGACA | TCT TGG CCA GCA TGG AAA ACT C |
| *hYAP* | TCCACCAGTGCAGCAGAATA | TTCCCATCCATCAGGAAGAG |
| *hWWTR1* | TCCCAGCCAAATCTCGTGATG | AGCGCATTGGGCATACTCAT |
| *hGATA3* | ATGCAAGTCCAGGC sCCAAG | TTGTGGTGGTCTGACAGTTCG |
| *hGATA2* | CCAGCTTCACCCCTAAGCAG | CCACAGTTGACACACTCCCG |
| *hTEAD1* | ATGGAAAGGATGAGTGACTCTGC | TCC CAC ATG GTG GAT AGA TAG C |
| *hSDC1* | CTATTCCCACGTCTCCAGAACC | CTATTCCCACGTCTCCAGAACC |
| *mMettl3* | CAAGCTGCACTTCAGACGAA | GCTTGGCGTGTGGTCTTT |
| *m18S* | AGTTCCAGCACATTTTGCGAG | TCATCCTCCGTGAGTTCTCCA |

**shRNA sequences:**

| | |
|---|---|
| *hMETTL3* | CGTCAGTATCTTGGGCAAGTT |
| | GCCTTAACATTGCCCACTGAT |
| | GCAAGTATGTTCACTATGAAA |
| *Scramble* | CCTAAGGTTAAGTCGCCCTCGC |
| *tetO-shMETTL3* | GCAAGTATGTTCACTATGAAA |
| *tetO-shGFP* | TACAACAGCCACAACGTCTAT |
| *mMettl3* | CGTCAGTATCTTGGGCAAATT |
| | CCTCAGTGGATCTGTTGTGAT |
| | GCACCCGCAAGATTGAGTTAT |

**Immunohistochemistry and Immunofluorescence assay:** Immunohistochemistry was performed using paraffin sections of human placenta. The slides were deparaffinized by histoclear and subsequently with 100%, 90%, 80% and 70% ethanol. Antigen retrieval was done using Decloaking chamber at 80°C for 15minutes. The slides were washed with 1X PBS and treated with 3% $H_2O_2$ to remove endogenous peroxidase followed by 3 times wash with 1X PBS. 10% goat serum was used as a blocking reagent for 1hour at RT followed by overnight incubation with 1:100 dilution of primary antibody or IgG at 4°C. These slides were washed 3 times with 1X PBS and incubated with biotin conjugated secondary antibody (1:200 dilution) for 1 hour at RT. The slides were washed with 1X PBS followed by treatment with streptavidin conjugated horseradish peroxidase (Vector laboratories, CA, SA-5704) for 20 minutes at RT. Reactivity was detected using DAB+ substrate chromogen system (Dako, TA-125-QHDX). The reaction was stopped in distilled water after sufficient color developed. The slides were counterstained with Mayer's hematoxylin for 2 minutes and washed with warm tap water until sufficient bluish coloration observed. The slides were then dehydrated by sequential treatment using 70%, 80%, 90%, 100% ethanol and histoclear. The sections were completely dried and mounted using Toluene as mountant and imaged using Nikon 90i microscope. Deparaffinized human placenta sections were rehydrated, and immunofluorescence was performed after antigen retrieval. The sections were washed with 0.25% Triton X-100 in 1X PBS and blocked for 1 hour using 10% fetal bovine serum (FBS). For mouse embryo cryosection, the sections were fixed using 4% paraformaldehyde in 1X PBS, permeabilized using 0.25% Triton X-100 in 1X PBS and blocked for 1 hour using 10% Normal Goat serum. The details of primary and secondary antibodies used are listed below. For mouse blastocyst, whole blastocysts were fixed using 4% paraformaldehyde in 1X PBS, permeabilized using 0.25% Triton X-100 in 1X PBS and blocked for 1 hour using 10% Normal Goat serum (Thermo Fisher scientific- 50062Z). The details of primary and secondary antibodies used are listed as supplementary materials.

**METTL3-fRIP:**

Formaldehyde crosslinking immunoprecipitation (fRIP) was processed as per the published protocol [1]. In brief, for fRIP sample preparation, proliferating hTSC were washed with PBS, crosslinked with 0.1%

formaldehyde for 10min at room temperature and crosslinking was stopped with glycine (150mM) for 10min at room temperature. Crosslinked cells were scrapped in 50ml of PBS, pelleted at 500g for 5 min at 4°C, snapped frozen in liquid nitrogen and stored at -80°C. Crosslinked cells were resuspended in IP buffer, sonicated the cell lysate using Bioruptor® Plus sonication. Cell lysates were cleared by centrifugation at 13000g for 15min at 4°C. Clear supernatant were again cleared using protein magnetic beads (washed with DEPC treated PBS) used to save for INPUT and immunoprecipitation using IgG and METTL3. Immunoprecipitated RNA were recovered following reverse crosslinking, phenol-chloroform-isoamyl alcohol extraction and precipitation using glycoblue. The IP RNA was purified using RNeasy Mini Kit (Qiagen-74104) using manufacturer's protocol where IP RNA were treated with DNaseI on the column. Purified IP RNA were used for stranded total RNASeq library preparation with ribodepletion and next-generation sequencing using NovaSeq 6000. Additional details of data analyses are mentioned in SI Appendix, Supplementary Materials and Methods.

**Collection and analyses of mouse embryos:** Preimplantation embryos were isolated at E3.5 matured blastocyst as described earlier [2]. E7.5 Embryos were used to test for METTL3 expression *via* immunostaining. The *Mettl3*$^{F/F}$ (female) and *Mettl3*$^{F/F}$*;Ubc-*$^{CreERT2}$ (male) animals were bred. Pregnant female animals were identified by presence of vaginal plug (gestational day 0.5) and embryos were harvested at e7.5 gestational days. Uterine horns from pregnant females were dissected out and separated EPC from rest of the embryos under microscope.
Embryo from each of the dissected conceptus were collected and genomic DNA preparation was done using Extract-N-Amp tissue PCR kit (Sigma- XNAT2). Respective primers are listed in the materials and methods section. Separated EPCs were placed in the center of 12 well plate coated with 0.1% gelatin and cultured like mTSC with 4-OH tamoxifen and assessed the growth of the explant as described earlier [3].
**Mouse TSC culture:** Mouse TSCs were cultured following previously described procedures [2]. Cells were harvested for total RNA extraction or fixed for immunofluorescence.

**RNA-seq analyses:**
Total RNA was used to construct RNA-seq libraries using the Illumina TruSeq Stranded Total RNA Sample Preparation Kit according to manufacturer's instructions. RNA seq was performed using Illumina HiSeq 2500 platform. The quality of the data sets was assessed using the FastQC v 0.11.9. In the next step mapping of the reads on reference genome was done using subread-align module of Subreads v 2.0.3 tool. The UCSC hg38 was utilized as reference downloaded from the iGenomes (https://support.illumina.com/ sequencing/sequencing_software/igenome.html). The subread-align was run using -t 0 (RNAseq data), -T 20 (number of processors), -a Gene.gtf along with --sortReadsByCoordinates argument while kept other arguments defaults. This produces the sorted bam file as output. We further index this bam file using the samtools index using argument -@ 20(number of processors). Along with it bamCoverage of deepTools v 3.5.0 was utilized to generate the bigWig file to visualized in IGV with the arguments -p 20 and –normalize using CPM. The sorted bam file of each of the sample was given input to the featureCounts of Subreads with the following arguments --countReadPairs -p -B -a Gene.gtf -T 20 -G genome.fa. A combined file is being generated to count each of the gene in each of samples. The meta file along with the count file generated from above step were given input to the DESeq2 v 1.34.0 module for finding the deferentially expressed genes. Heatmap were generated using pheatmap () function of r package pheatmap v1.0.12. PCA plot were obtained using plotPCA() function of DESeq2. MA plot was generated using ggmaplot () function of r package ggpubr v0.4.0. Enhanced Volcano () function of bio-conductor package EnhancedVolcano v 1.12.0 was utilized in default mode to generate the Volcano plot.

**m6A RNA Cut&Run data analyses:**
Single-end 50-bp stranded libraries were sequenced using HiSeq2500. Multiplexed reads were split based on their barcodes using Illumina Basespace. Reads were trimmed to remove the TruSeq adaptor using trim_galore with parameters '-q 0 -a AGATCGGAAGAGCACACGTCTGAACTCCAGTCAC–phred33–fastqc'. The mapped reads were used to call peaks following "narrowPeak" file with *p*-value 1e-$^{05}$ which is output of the macs2 was processed using "cut -f 1-6" to convert into bed file which were used as a input file for peaks annotation and motif finding.

**METTL3-fRIP data analyses:**

Paired-end 100-bp stranded libraries were sequenced using HiSeq2500. The mapped reads were used to call peaks following "narrowPeak" file with *p*-value 1e-$^{05}$ which is output of the macs2 was processed using "cut -f 1-6" to convert into bed file which were used as an input file for peaks annotation and motif finding.

**Distribution of peaks in different genomics region:**
To get the genomics distribution of peaks the bed file was imported using toGRanges() function with specification of format as "BED", which convert the coordinate into genomics ranges. In the next step annotatePeak() function of ChIPpeakAnno Bioconductor package was called to annotate the peaks available in genomics ranges with reference annotation database "org.Hs.eg.db". Finally the function plotAnnoPie() function was used to plot pie-chart of genomics distribution of peaks.

**Differential m6A peak identification:**
To identifying m6A peaks that are differentially enriched between the control and METTL3KD samples, we used an R Bioconductor package DiffBind[4] (Available online at: http://bioconductor.org/packages/release/bioc/html/DiffBind.html). MACS2 generated m6A peaks (*p*-value 1e-$^{05}$) from GFPKD and METTL3KD were used to generate Intersection of genomic regions (peaks) using BedSect[5]. Intersection of m6A peaks and Bam files GFPKD and METTL3KD were used for this purpose. Volcano plots were generated using the default DESeq2 analysis parameters (see Section 10.3 discussing the technical details of the analysis, default threshold of FDR <= 0.05).

**Differential m6A peak associated gene identification and functional association:**
Reproducible differentially enriched peaks associated coordinates were used to identify m6A associated genes using GREAT tool (GREAT version 4.0.4)[6]. We also utilized this tool to get functional association and pathways information associated m6A genes. We also used EnrichR[7] and PlacentaCellEnrich[8] for functional association and pathways. Downstream analysis for the pathways and association with different pathological conditions were visualized using DeepVenn.

**Preparation of coverage file for visualization:**
Here first of all reads were filtered using "samtools view" of samtools v 1.6[9] with argument -F 516 and -q 30. The output bam file was then sorted using "samtools sort" with parameter -@ 20. The resulted bam is then given input to "samtools rmdup" to remove the duplicated reads. Next these bam files were indexed using "samtools index" with parameter "-@ 20". Finally bamCoverage v 3.5.1 of deepTools[10] was used with all most all default parameters except those are "-p 20", " -effective Genome Size 2913022398", "--normalize Using BPM" to get the final coverage file in the form of Bigwig files for visualization using IGV tool[11].

**Peak annotation and motif identification:**
The peaks files were further filtered using "bedtools intersect" function with hg38 blacklisted regions. The curated bed file from this step was given input to the "annotatePeaks.pl" of HOMER[12] for annotation of the peaks. In the next step sequence of each of the peaks was extracted using "bedtools get fasta" function where the UCSC hg38 obtained from iGenomics was used as reference sequence. The motif was identified using "findMotifs.pl" script of HOMER with the parameters "-len 5,6,7,8", "-rna", "-p 20", "-S 50" and "-mask".

**Cut&Run and Data Analyses**
Proliferating semiconfluent 200,000 live hTSC were used per sample for Cut&Run following published protocol[13,14]. Trypsinized hTSC were washed twice with the wash buffer (Wash buffer was prepared by adding 1ml 1M HEPES pH7.5, 1.5ml 5M NaCl, 12.5µL 2M Spermidine and 1 Roche Complete Protease Inhibitor EDTA-Free tablet to final volume to 50ml with dH2O). Concanavalin A-coated beads (10µL/sample from EpiCypher) were washed with binding buffer (binding buffer was prepared by mixing 400µL 1M HEPES-KOH pH 7.9, 200µL 1M KCl, 20µL 1M $CaCl_2$ and 20µL 1M $MnCl_2$, and bring the final volume to 20 ml with $dH_2O$. Stored up to 6 months at 4°C.) and recovered by placing on the magnetic stand. Washed Concanavalin-A coated beads were resuspended in wash buffer and incubated with the hTSC under rotation at room temperature for 10minutes. Using a magnetic stand, the cells bound to the Concanavalin A-coated beads were separated from the suspension, the liquid was discarded, and the

cells were resuspended in 100µl of antibody buffer (antibody buffer was prepared by adding 8µL 0.5 M EDTA with 2 ml Dig-wash buffer) containing 0.5% (wt/vol) Digitonin (Dig-wash buffer). Primary antibody of interest (METTL3 or IgG antibody; 1:100 dilution) was added to the samples and incubated overnight at 4°C with intermittent shaking. The samples were washed twice with 150µL of Dig-wash buffer (200µL 5% Digitonin with 20ml Wash buffer) while keeping the samples on ice, magnetic stand was used to remove the residual liquid. Next, these cells bound to the Concanavalin A-coated beads were resuspended in 50µl of Dig-wash buffer and incubated with 1.5µl A/G-MNase (from Cell Signaling) for 1 hour with intermittent shaking/rotation in cold room. Incubation was followed by vortexing, a quick spin and then placed on the magnet stand to separate out and discard the liquid. The samples were washed twice with Dig-wash buffer and resuspended in the 100µL of Dig-wash buffer and placed on the metal eppendorf rack equilibrated in an ice-water bath. A/G-MNase reaction was activated by adding 2µL 100 mM $CaCl_2$ to each sample, mixed well and incubated the tubes for 30 minutes on the metal rack placed in an ice-water bath. The MNase reaction was stopped by adding equal volume of 2X STOP buffer (stop buffer was prepared by adding 3.44ml $dH_2O$, 136µl 5M NaCl, 400µL 0.2M EGTA, 40µL 5% Digitonin, 10µL RNase A, 20µL 20 mg/ml glycogen). These samples were mixed by vortexing and incubated for 10 minutes at 37°C to release CUT&RUN fragments from the insoluble nuclear chromatin. Release CUT&RUN fragments were recovered by centrifugation for 5 min in 4°C at 16,000g and placed on magnetic stand. Supernatant with CUT&RUN fragments were transferred into a fresh 1.5ml microcentrifuge tube containing 2µL of 10% SDS and 2.5µL of Proteinase K (Thermo Scientific). These samples were incubated for 10 minutes at 70°C. CUT&RUN DNA fragments were purified by using a phase-lock tube (Qiagen) and Phenol–chloroform–isoamyl alcohol. The purified CUT&RUN DNA fragments were transferred to a fresh tube and precipitated following ethanol precipitation. The dried CUT&RUN DNA pellet was dissolved in 20µL 1 mM Tris-HCl pH8 0.1 mM EDTA. The DNA was quantified using 1µL of the freshly dissolved pellet, for example using fluorescence detection with a Qubit instrument (Life Technologies, Waltham, MA). Evaluated the presence of cleaved fragments and the size distribution by capillary electrophoresis with fluorescence detection using a 4200 TapeStation System (Agilent). The library was prepared following protocol from Swift bioscience 1S Plus Combinatorial Dual Indexing Kit and Accel-NGS 1S Plus DNA Library Kit. We have used “Chipsequer nf-core” pipeline (3) for downstream analyses of CUT&RUN data. Reproducible METTL3 associated peak coordinates were used to identify METTL3 regulated genes using GREAT tool (GREAT version 4.0.4)[6].
