## Supplementary material for "METTL3 shapes m6A epitranscriptomic landscape for successful human placentation": S4

### Homer *de novo* Motif Results (Homer_motifResults//)

Known Motif Enrichment Results
Gene Ontology Enrichment Results
If Homer is having trouble matching a motif to a known motif, try copy/pasting the matrix file into STAMP
More information on motif finding results: HOMER | Description of Results | Tips
Total target sequences = 8008
Total background sequences = 32820
* - possible false positive

| Rank | Motif | P-value | log P-pvalue | % of Targets | % of Background | STD(Bg STD) | Best Match/Details | Motif File |
|---|---|---|---|---|---|---|---|---|
| 1 | 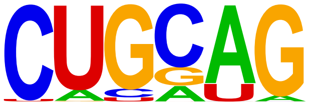  | 1e-209 | -4.817e+02 | 43.97% | 25.93% | 72.7bp (72.4bp) | hsa-miR-4418 MIMAT0018930 Homo sapiens miR-4418 Targets (miRBase)(0.807) More Information \| Similar Motifs Found | motif file (matrix) |
| 2 | 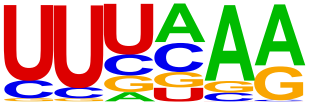  | 1e-125 | -2.884e+02 | 48.09% | 33.60% | 72.5bp (65.3bp) | hsa-miR-1305 MIMAT0005893 Homo sapiens miR-1305 Targets (miRBase)(0.684) More Information \| Similar Motifs Found | motif file (matrix) |
| 3 | 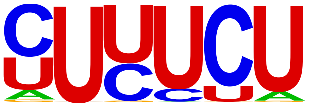  | 1e-99 | -2.280e+02 | 82.59% | 71.34% | 100.3bp (177.3bp) | hsa-miR-3160-5p MIMAT0019212 Homo sapiens miR-3160-5p Targets (miRBase)(0.738) More Information \| Similar Motifs Found | motif file (matrix) |
| 4 | 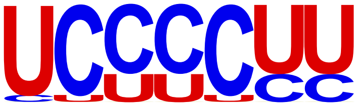  | 1e-98 | -2.262e+02 | 73.44% | 61.01% | 136.4bp (235.8bp) | hsa-miR-3127-3p MIMAT0019201 Homo sapiens miR-3127-3p Targets (miRBase)(0.730) More Information \| Similar Motifs Found | motif file (matrix) |
| 5 | 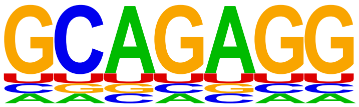  | 1e-90 | -2.084e+02 | 88.37% | 78.89% | 0bp (0bp) | hsa-miR-2467-3p MIMAT0019953 Homo sapiens miR-2467-3p Targets (miRBase)(0.862) More Information \| Similar Motifs | motif file (matrix) |

| | | | | | | | | |
|---|---|---|---|---|---|---|---|---|
| | | | | | | | Found | |
| 6 | 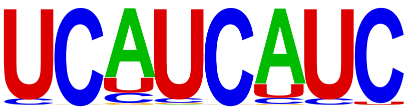 | 1e-78 | -1.814e+02 | 5.81% | 1.72% | 72.0bp (117.1bp) | hsa-miR-1272 MIMAT0005925 Homo sapiens miR-1272 Targets (miRBase)(0.809) More Information \| Similar Motifs Found | motif file (matrix) |
| 7 | 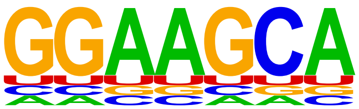 | 1e-74 | -1.719e+02 | 89.79% | 81.65% | 0bp (0bp) | hsa-miR-516a-3p MIMAT0006778 Homo sapiens miR-516a-3p Targets (miRBase)(0.817) More Information \| Similar Motifs Found | motif file (matrix) |
| 8 | 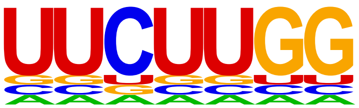 | 1e-69 | -1.594e+02 | 93.06% | 86.21% | 0bp (0bp) | hsa-miR-578 MIMAT0003243 Homo sapiens miR-578 Targets (miRBase) (0.704) More Information \| Similar Motifs Found | motif file (matrix) |
| 9 | 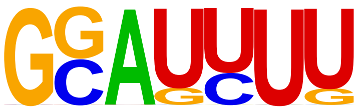 | 1e-59 | -1.361e+02 | 65.26% | 55.30% | 59.0bp (63.9bp) | hsa-miR-1290 MIMAT0005880 Homo sapiens miR-1290 Targets (miRBase)(0.821) More Information \| Similar Motifs Found | motif file (matrix) |
| 10 | 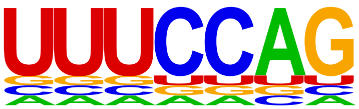 | 1e-41 | -9.601e+01 | 94.58% | 89.96% | 0bp (0bp) | hsa-miR-875-3p MIMAT0004923 Homo sapiens miR-875-3p Targets (miRBase)(0.870) More Information \| Similar Motifs Found | motif file (matrix) |
| 11 | 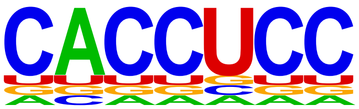 | 1e-39 | -9.123e+01 | 81.06% | 74.10% | 0bp (0bp) | hsa-miR-3689d MIMAT0019008 Homo sapiens miR-3689d Targets (miRBase)(0.862) More Information \| Similar Motifs Found | motif file (matrix) |
| 12 | 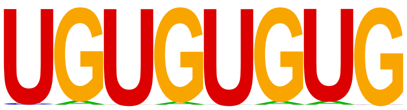 | 1e-30 | -7.008e+01 | 1.56% | 0.33% | 64.9bp (59.3bp) | hsa-miR-147 MIMAT0000251 Homo sapiens miR-147 Targets (miRBase) (0.751) More Information \| Similar Motifs Found | motif file (matrix) |

| | | | | | | | | |
|---|---|---|---|---|---|---|---|---|
| 13 | 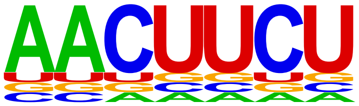 | 1e-29 | -6.898e+01 | 93.49% | 89.43% | 0bp (0bp) | hsa-miR-3653 MIMAT0018073 Homo sapiens miR-3653 Targets (miRBase)(0.740) More Information \| Similar Motifs Found | motif file (matrix) |
| 14 | 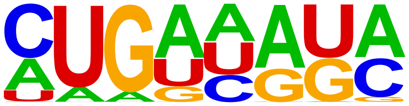 | 1e-24 | -5.580e+01 | 25.84% | 20.49% | 60.8bp (53.7bp) | hsa-miR-205* MIMAT0009197 Homo sapiens miR-205* Targets (miRBase)(0.699) More Information \| Similar Motifs Found | motif file (matrix) |
| 15 | 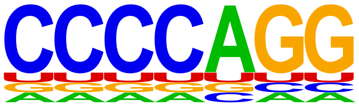 | 1e-23 | -5.390e+01 | 86.65% | 82.04% | 0bp (0bp) | hsa-miR-1915 MIMAT0007892 Homo sapiens miR-1915 Targets (miRBase)(0.802) More Information \| Similar Motifs Found | motif file (matrix) |
| 16 | 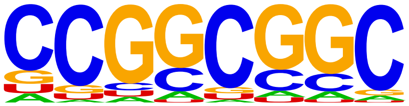 | 1e-22 | -5.207e+01 | 2.21% | 0.81% | 57.3bp (45.2bp) | hsa-miR-3960 MIMAT0019337 Homo sapiens miR-3960 Targets (miRBase)(0.710) More Information \| Similar Motifs Found | motif file (matrix) |
| 17 | 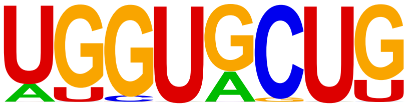 | 1e-21 | -4.908e+01 | 2.95% | 1.31% | 55.1bp (49.4bp) | hsa-miR-3065-3p MIMAT0015378 Homo sapiens miR-3065-3p Targets (miRBase)(0.826) More Information \| Similar Motifs Found | motif file (matrix) |
| 18 | 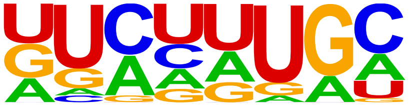 | 1e-18 | -4.208e+01 | 74.04% | 69.04% | 86.3bp (66.7bp) | hsa-miR-502-5p MIMAT0002873 Homo sapiens miR-502-5p Targets (miRBase)(0.663) More Information \| Similar Motifs Found | motif file (matrix) |
| 19 * | 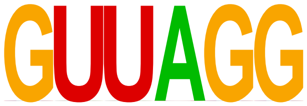 | 1e-11 | -2.726e+01 | 97.70% | 96.17% | 72.7bp (64.0bp) | hsa-miR-1258 MIMAT0005909 Homo sapiens miR-1258 Targets (miRBase)(0.805) More Information \| Similar Motifs Found | motif file (matrix) |

| | | | | | | | | |
|---|---|---|---|---|---|---|---|---|
| 20 * | 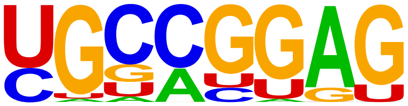 | 1e-10 | -2.417e+01 | 83.74% | 80.60% | 88.0bp (65.9bp) | hsa-miR-1976 MIMAT0009451 Homo sapiens miR-1976 Targets (miRBase)(0.703) More Information \| Similar Motifs Found | motif file (matrix) |
| 21 * | 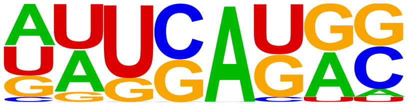 | 1e-9 | -2.101e+01 | 62.81% | 59.13% | 71.1bp (98.1bp) | hsa-miR-487a MIMAT0002178 Homo sapiens miR-487a Targets (miRBase)(0.636) More Information \| Similar Motifs Found | motif file (matrix) |
| 22 * | 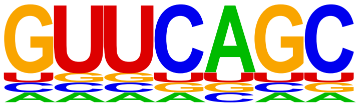 | 1e-7 | -1.817e+01 | 95.79% | 94.25% | 0bp (0bp) | hsa-miR-4539 MIMAT0019082 Homo sapiens miR-4539 Targets (miRBase)(0.788) More Information \| Similar Motifs Found | motif file (matrix) |
| 23 * | 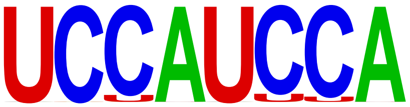 | 1e-6 | -1.420e+01 | 0.57% | 0.22% | 57.4bp (51.5bp) | hsa-miR-432* MIMAT0002815 Homo sapiens miR-432* Targets (miRBase)(0.792) More Information \| Similar Motifs Found | motif file (matrix) |
| 24 * | 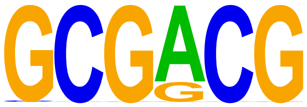 | 1e-3 | -7.638e+00 | 1.47% | 1.02% | 58.9bp (44.2bp) | hsa-miR-1181 MIMAT0005826 Homo sapiens miR-1181 Targets (miRBase)(0.798) More Information \| Similar Motifs Found | motif file (matrix) |
| 25 * | 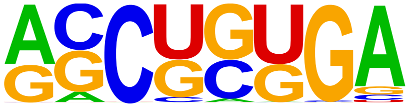 | 1e-3 | -7.104e+00 | 2.97% | 2.35% | 54.2bp (42.4bp) | hsa-miR-513a-5p MIMAT0002877 Homo sapiens miR-513a-5p Targets (miRBase)(0.708) More Information \| Similar Motifs Found | motif file (matrix) |
| 26 * | 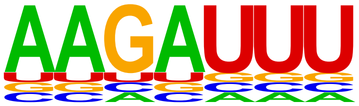 | 1e-2 | -6.878e+00 | 95.10% | 94.23% | 0bp (0bp) | hsa-miR-4762-5p MIMAT0019910 Homo sapiens miR-4762-5p Targets (miRBase)(0.795) More Information \| Similar Motifs Found | motif file (matrix) |

| | | | | | | | | |
|---|---|---|---|---|---|---|---|---|
| 27 * | 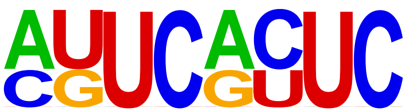 | 1e-2 | -6.588e+00 | 0.86% | 0.55% | 118.3bp (50.7bp) | hsa-miR-668 MIMAT0003881 Homo sapiens miR-668 Targets (miRBase) (0.694) More Information \| Similar Motifs Found | motif file (matrix) |
| 28 * | 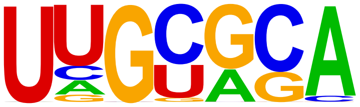 | 1e-2 | -6.267e+00 | 23.34% | 21.83% | 74.0bp (56.0bp) | hsa-let-7i* MIMAT0004585 Homo sapiens let-7i* Targets (miRBase) (0.796) More Information \| Similar Motifs Found | motif file (matrix) |
| 29 * | 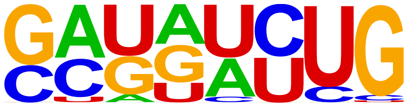 | 1e-1 | -3.865e+00 | 9.25% | 8.53% | 49.6bp (63.2bp) | hsa-miR-4748 MIMAT0019884 Homo sapiens miR-4748 Targets (miRBase)(0.620) More Information \| Similar Motifs Found | motif file (matrix) |
| 30 * | 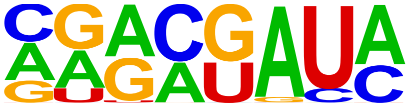 | 1e0 | -1.106e+00 | 42.84% | 42.57% | 95.5bp (64.8bp) | hsa-miR-625* MIMAT0004808 Homo sapiens miR-625* Targets (miRBase)(0.615) More Information \| Similar Motifs Found | motif file (matrix) |
| 31 * | 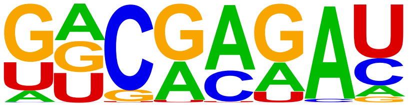 | 1e0 | -4.182e-01 | 0.90% | 0.94% | 59.6bp (64.9bp) | hsa-miR-31 MIMAT0000089 Homo sapiens miR-31 Targets (miRBase) (0.682) More Information \| Similar Motifs Found | motif file (matrix) |
| 32 * | 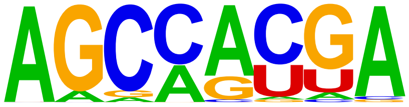 | 1e0 | -2.915e-01 | 6.12% | 6.31% | 88.9bp (70.0bp) | hsa-miR-4474-3p MIMAT0019001 Homo sapiens miR-4474-3p Targets (miRBase)(0.662) More Information \| Similar Motifs Found | motif file (matrix) |
| 33 * | 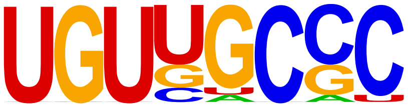 | 1e0 | -1.373e-02 | 0.75% | 1.00% | 29.7bp (16.0bp) | hsa-miR-196a* MIMAT0004562 Homo sapiens miR-196a* Targets (miRBase)(0.748) More Information \| Similar Motifs Found | motif file (matrix) |

| | | | | | | | | |
|---|---|---|---|---|---|---|---|---|
| 34 * | 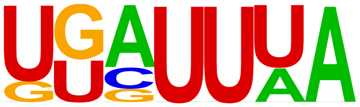 | 1e0 | 0.000e+00 | 100.00% | 100.00% | 78.3bp (63.8bp) | hsa-miR-4503 MIMAT0019039 Homo sapiens miR-4503 Targets (miRBase)(0.641) More Information \| Similar Motifs Found | motif file (matrix) |
