## Supplementary material for "METTL3 shapes m6A epitranscriptomic landscape for successful human placentation": S5

### Homer *de novo* Motif Results (Homer_motifResults//)

Known Motif Enrichment Results
Gene Ontology Enrichment Results
If Homer is having trouble matching a motif to a known motif, try copy/pasting the matrix file into STAMP
More information on motif finding results: HOMER | Description of Results | Tips
Total target sequences = 2363
Total background sequences = 9519
* - possible false positive

| Rank | Motif | P-value | log P-pvalue | % of Targets | % of Background | STD(Bg STD) | Best Match/Details | Motif File |
|---|---|---|---|---|---|---|---|---|
| 1 | 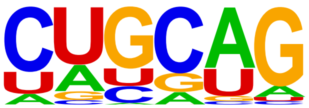 | 1e-70 | -1.630e+02 | 59.97% | 39.54% | 81.7bp (296.5bp) | hsa-miR-4418 MIMAT0018930 Homo sapiens miR-4418 Targets (miRBase) (0.781) More Information \| Similar Motifs Found | motif file (matrix) |
| 2 | 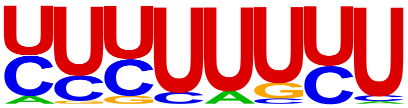 | 1e-57 | -1.333e+02 | 78.29% | 61.09% | 70.6bp (666.6bp) | hsa-miR-3613-3p MIMAT0017991 Homo sapiens miR-3613-3p Targets (miRBase)(0.723) More Information \| Similar Motifs Found | motif file (matrix) |
| 3 | 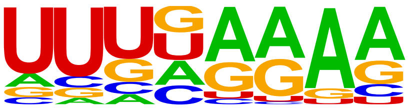 | 1e-52 | -1.205e+02 | 19.76% | 8.19% | 49.9bp (58.4bp) | hsa-miR-3658 MIMAT0018078 Homo sapiens miR-3658 Targets (miRBase) (0.670) More Information \| Similar Motifs Found | motif file (matrix) |
| 4 | 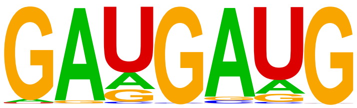 | 1e-47 | -1.105e+02 | 12.48% | 3.94% | 55.5bp (86.0bp) | hsa-miR-1322 MIMAT0005953 Homo sapiens miR-1322 Targets (miRBase) (0.788) More Information \| Similar Motifs Found | motif file (matrix) |
| 5 | 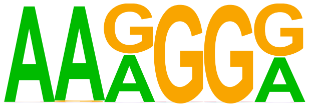 | 1e-46 | -1.067e+02 | 74.02% | 58.26% | 71.2bp (94.8bp) | hsa-miR-4287 MIMAT0016917 Homo sapiens miR-4287 Targets (miRBase) (0.717) More Information \| Similar Motifs | motif file (matrix) |

| | | | | | | | | |
|---|---|---|---|---|---|---|---|---|
| | | | | | | | Found | |
| 6 | 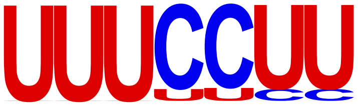  | 1e-43 | -1.009e+02 | 78.25% | 63.51% | 77.4bp (772.1bp) | hsa-miR-1252 MIMAT0005944 Homo sapiens miR-1252 Targets (miRBase) (0.774) More Information \| Similar Motifs Found | motif file (matrix) |
| 7 | 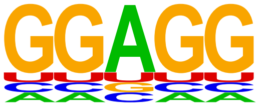  | 1e-40 | -9.306e+01 | 92.26% | 81.65% | 68.2bp (106.3bp) | hsa-miR-4266 MIMAT0016892 Homo sapiens miR-4266 Targets (miRBase) (0.769) More Information \| Similar Motifs Found | motif file (matrix) |
| 8 | 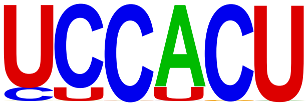  | 1e-35 | -8.263e+01 | 77.87% | 64.62% | 72.0bp (761.8bp) | hsa-miR-3681 MIMAT0018108 Homo sapiens miR-3681 Targets (miRBase) (0.790) More Information \| Similar Motifs Found | motif file (matrix) |
| 9 | 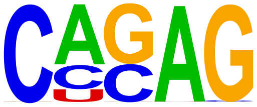  | 1e-28 | -6.465e+01 | 42.62% | 30.46% | 65.1bp (66.4bp) | hsa-miR-4257 MIMAT0016878 Homo sapiens miR-4257 Targets (miRBase) (0.704) More Information \| Similar Motifs Found | motif file (matrix) |
| 10 | 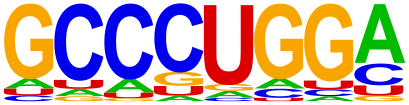  | 1e-13 | -3.111e+01 | 6.18% | 2.78% | 46.6bp (82.4bp) | hsa-miR-4308 MIMAT0016861 Homo sapiens miR-4308 Targets (miRBase) (0.805) More Information \| Similar Motifs Found | motif file (matrix) |
| 11 | 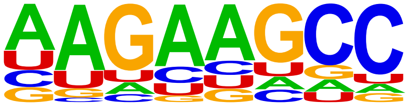  | 1e-12 | -2.869e+01 | 8.25% | 4.38% | 41.1bp (42.4bp) | hsa-miR-103a-2* MIMAT0009196 Homo sapiens miR-103a-2* Targets (miRBase)(0.738) More Information \| Similar Motifs Found | motif file (matrix) |
| 12 * | 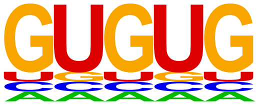  | 1e-11 | -2.601e+01 | 96.91% | 93.45% | 73.5bp (266.9bp) | hsa-miR-4455 MIMAT0018977 Homo sapiens miR-4455 Targets (miRBase) (0.762) More Information \| Similar Motifs Found | motif file (matrix) |

| | | | | | | | | |
|---|---|---|---|---|---|---|---|---|
| 13 * | 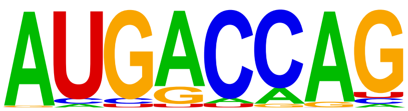 | 1e-9 | -2.234e+01 | 4.23% | 1.87% | 56.0bp (48.6bp) | hsa-miR-4435 MIMAT0018951 Homo sapiens miR-4435 Targets (miRBase) (0.723) More Information \| Similar Motifs Found | motif file (matrix) |
| 14 * | 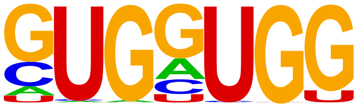 | 1e-7 | -1.790e+01 | 7.49% | 4.54% | 44.6bp (70.8bp) | hsa-miR-4456 MIMAT0018978 Homo sapiens miR-4456 Targets (miRBase) (0.789) More Information \| Similar Motifs Found | motif file (matrix) |
| 15 * | 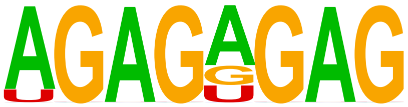 | 1e0 | -8.604e-01 | 4.99% | 4.88% | 44.0bp (104.2bp) | hsa-miR-4469 MIMAT0018996 Homo sapiens miR-4469 Targets (miRBase) (0.772) More Information \| Similar Motifs Found | motif file (matrix) |
| 16 * | 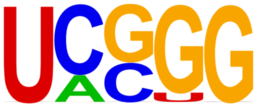 | 1e0 | -5.782e-01 | 24.38% | 24.51% | 62.9bp (547.0bp) | hsa-miR-3181 MIMAT0015061 Homo sapiens miR-3181 Targets (miRBase) (0.700) More Information \| Similar Motifs Found | motif file (matrix) |
| 17 * | 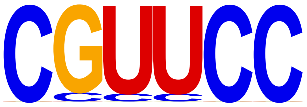 | 1e0 | -0.000e+00 | 13.88% | 18.31% | 65.5bp (475.7bp) | hsa-miR-1292 MIMAT0005943 Homo sapiens miR-1292 Targets (miRBase) (0.776) More Information \| Similar Motifs Found | motif file (matrix) |
