## Supplementary material for "METTL3 shapes m6A epitranscriptomic landscape for successful human placentation": S8

### Homer *de novo* Motif Results (Homer_motifResults//)

Known Motif Enrichment Results
Gene Ontology Enrichment Results
If Homer is having trouble matching a motif to a known motif, try copy/pasting the matrix file into STAMP
More information on motif finding results: HOMER | Description of Results | Tips
Total target sequences = 16986
Total background sequences = 70166
* - possible false positive

| Rank | Motif | P-value | log P-pvalue | % of Targets | % of Background | STD(Bg STD) | Best Match/Details | Motif File |
|---|---|---|---|---|---|---|---|---|
| 1 | 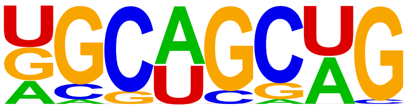 | 1e-1322 | -3.046e+03 | 49.68% | 19.24% | 602.8bp (415.9bp) | hsa-miR-5047 MIMAT0020541 Homo sapiens miR-5047 Targets (miRBase)(0.727) More Information \| Similar Motifs Found | motif file (matrix) |
| 2 | 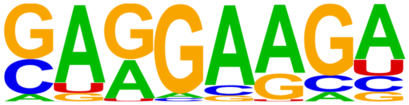 | 1e-983 | -2.264e+03 | 61.01% | 32.71% | 561.0bp (401.6bp) | hsa-miR-4778-3p MIMAT0019937 Homo sapiens miR-4778-3p Targets (miRBase)(0.766) More Information \| Similar Motifs Found | motif file (matrix) |
| 3 | 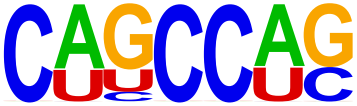 | 1e-954 | -2.199e+03 | 65.43% | 37.33% | 589.5bp (412.8bp) | hsa-miR-3064-5p MIMAT0019864 Homo sapiens miR-3064-5p Targets (miRBase)(0.800) More Information \| Similar Motifs Found | motif file (matrix) |
| 4 | 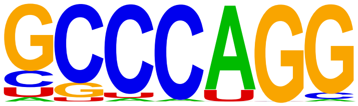 | 1e-913 | -2.103e+03 | 43.00% | 18.40% | 618.5bp (424.0bp) | hsa-miR-4292 MIMAT0016919 Homo sapiens miR-4292 Targets (miRBase)(0.827) More Information \| Similar Motifs Found | motif file (matrix) |
| 5 | 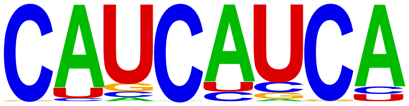 | 1e-904 | -2.082e+03 | 74.85% | 47.94% | 569.6bp (414.0bp) | hsa-miR-4502 MIMAT0019038 Homo sapiens miR-4502 Targets (miRBase)(0.767) More Information \| Similar Motifs | motif file (matrix) |

| | | | | | | | | |
|---|---|---|---|---|---|---|---|---|
| | | | | | | | Found | |
| 6 | 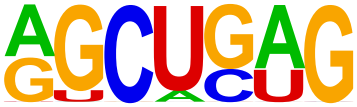 | 1e-804 | -1.851e+03 | 64.78% | 38.96% | 584.9bp (422.1bp) | hsa-miR-4663 MIMAT0019735 Homo sapiens miR-4663 Targets (miRBase)(0.740) More Information \| Similar Motifs Found | motif file (matrix) |
| 7 | 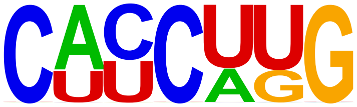 | 1e-741 | -1.708e+03 | 59.61% | 34.94% | 602.3bp (417.7bp) | hsa-miR-4761-5p MIMAT0019908 Homo sapiens miR-4761-5p Targets (miRBase)(0.783) More Information \| Similar Motifs Found | motif file (matrix) |
| 8 | 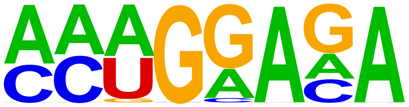 | 1e-688 | -1.586e+03 | 48.42% | 25.69% | 577.1bp (395.8bp) | hsa-miR-4768-3p MIMAT0019921 Homo sapiens miR-4768-3p Targets (miRBase)(0.723) More Information \| Similar Motifs Found | motif file (matrix) |
| 9 | 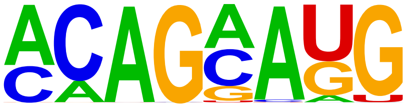 | 1e-609 | -1.405e+03 | 44.21% | 23.32% | 536.2bp (403.5bp) | hsa-miR-214 MIMAT0000271 Homo sapiens miR-214 Targets (miRBase) (0.729) More Information \| Similar Motifs Found | motif file (matrix) |
| 10 | 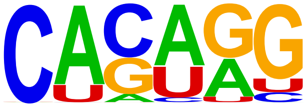 | 1e-584 | -1.345e+03 | 62.88% | 40.84% | 580.8bp (413.0bp) | hsa-miR-513a-5p MIMAT0002877 Homo sapiens miR-513a-5p Targets (miRBase)(0.736) More Information \| Similar Motifs Found | motif file (matrix) |
| 11 | 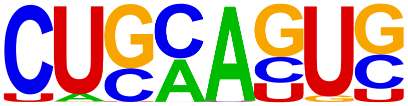 | 1e-523 | -1.205e+03 | 50.17% | 29.93% | 596.9bp (416.2bp) | hsa-miR-4418 MIMAT0018930 Homo sapiens miR-4418 Targets (miRBase)(0.816) More Information \| Similar Motifs Found | motif file (matrix) |
| 12 | 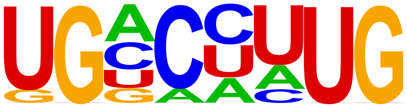 | 1e-513 | -1.182e+03 | 49.96% | 29.93% | 576.2bp (411.1bp) | hsa-miR-3922-5p MIMAT0019227 Homo sapiens miR-3922-5p Targets (miRBase)(0.694) More Information \| Similar Motifs Found | motif file (matrix) |

| | | | | | | | | |
|---|---|---|---|---|---|---|---|---|
| 13 | 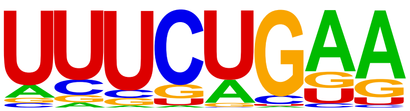 | 1e-468 | -1.080e+03 | 45.41% | 26.70% | 543.6bp (393.1bp) | hsa-miR-3942-3p MIMAT0019230 Homo sapiens miR-3942-3p Targets (miRBase)(0.688) More Information \| Similar Motifs Found | motif file (matrix) |
| 14 | 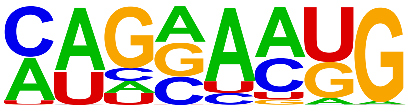 | 1e-422 | -9.730e+02 | 27.06% | 12.65% | 526.6bp (418.0bp) | hsa-miR-4277 MIMAT0016908 Homo sapiens miR-4277 Targets (miRBase)(0.680) More Information \| Similar Motifs Found | motif file (matrix) |
| 15 | 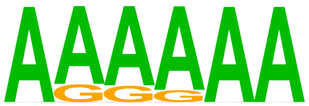 | 1e-388 | -8.952e+02 | 64.64% | 46.70% | 551.7bp (382.8bp) | hsa-miR-3613-3p MIMAT0017991 Homo sapiens miR-3613-3p Targets (miRBase)(0.774) More Information \| Similar Motifs Found | motif file (matrix) |
| 16 | 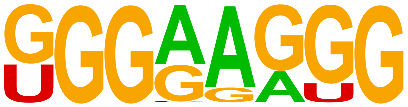 | 1e-388 | -8.940e+02 | 26.62% | 12.81% | 565.5bp (376.5bp) | hsa-miR-3202 MIMAT0015089 Homo sapiens miR-3202 Targets (miRBase)(0.797) More Information \| Similar Motifs Found | motif file (matrix) |
| 17 | 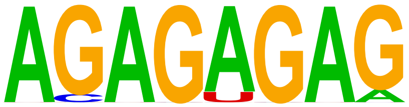 | 1e-332 | -7.650e+02 | 39.90% | 24.58% | 555.8bp (391.6bp) | hsa-miR-4311 MIMAT0016863 Homo sapiens miR-4311 Targets (miRBase)(0.735) More Information \| Similar Motifs Found | motif file (matrix) |
| 18 | 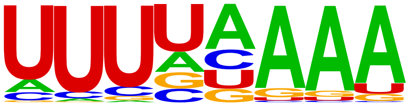 | 1e-329 | -7.585e+02 | 24.51% | 12.14% | 529.6bp (365.4bp) | hsa-miR-450b-5p MIMAT0004909 Homo sapiens miR-450b-5p Targets (miRBase)(0.621) More Information \| Similar Motifs Found | motif file (matrix) |
| 19 | 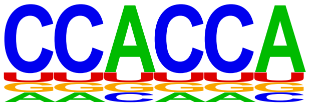 | 1e-286 | -6.590e+02 | 23.27% | 11.91% | 628.8bp (412.7bp) | hsa-miR-4456 MIMAT0018978 Homo sapiens miR-4456 Targets (miRBase)(0.835) More Information \| Similar Motifs Found | motif file (matrix) |

| | | | | | | | | |
|---|---|---|---|---|---|---|---|---|
| 20 | 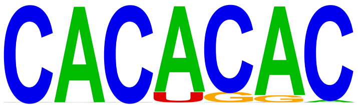 | 1e-189 | -4.368e+02 | 15.49% | 7.75% | 615.2bp (392.3bp) | hsa-miR-147 MIMAT0000251 Homo sapiens miR-147 Targets (miRBase) (0.797) More Information \| Similar Motifs Found | motif file (matrix) |
| 21 | 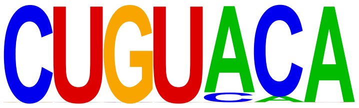 | 1e-172 | -3.970e+02 | 22.39% | 13.43% | 533.4bp (415.3bp) | hsa-let-7g* MIMAT0004584 Homo sapiens let-7g* Targets (miRBase) (0.868) More Information \| Similar Motifs Found | motif file (matrix) |
| 22 | 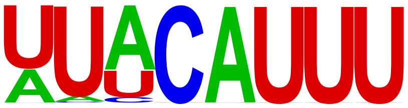 | 1e-135 | -3.129e+02 | 19.99% | 12.36% | 525.6bp (390.0bp) | hsa-miR-3646 MIMAT0018065 Homo sapiens miR-3646 Targets (miRBase)(0.781) More Information \| Similar Motifs Found | motif file (matrix) |
| 23 | 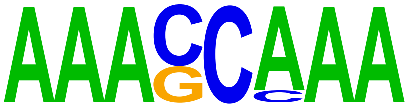 | 1e-82 | -1.909e+02 | 6.38% | 3.04% | 555.7bp (380.6bp) | hsa-miR-9 MIMAT0000441 Homo sapiens miR-9 Targets (miRBase) (0.681) More Information \| Similar Motifs Found | motif file (matrix) |
| 24 | 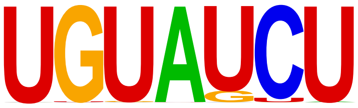 | 1e-43 | -9.985e+01 | 10.01% | 6.79% | 587.0bp (396.5bp) | hsa-miR-297 MIMAT0004450 Homo sapiens miR-297 Targets (miRBase) (0.704) More Information \| Similar Motifs Found | motif file (matrix) |
